# Mutation bias alters the distribution of fitness effects of mutations

**DOI:** 10.1101/2024.03.24.586369

**Authors:** Mrudula Sane, Shazia Parveen, Deepa Agashe

## Abstract

Mutation bias is an important factor determining the diversity of genetic variants available for selection. As adaptation proceeds and some beneficial mutations are fixed, new beneficial mutations become rare, limiting further adaptation. The depletion of beneficial mutations is especially stark within the mutational class favoured by the existing mutation bias. Recent theoretical work predicts that this problem may be alleviated by a change in the direction of mutation bias (i.e., a bias reversal). If populations sample previously underexplored types of mutations, the distribution of fitness effects (DFE) of mutations should shift towards more beneficial mutations. Here, we test this prediction using *Escherichia coli*, which has a transition mutation bias, with ∼54% single-nucleotide mutations being transitions compared to the unbiased expectation of ∼33% transitions. We generated mutant strains with a wide range of mutation biases, from 97% transitions to 98% transversions, either reinforcing or reversing the wild type transition bias. Quantifying DFEs of ∼100 single mutations obtained from mutation accumulation experiments for each strain, we find strong support for the theoretical prediction. Strains that oppose the ancestral bias (i.e., with a strong transversion bias) have DFEs with the highest proportion of beneficial mutations, whereas strains that exacerbate the ancestral transition bias have up to 10-fold fewer beneficial mutations. Such dramatic differences in the DFE should drive large variation in the rate and outcome of adaptation, suggesting an important and generalized evolutionary role for mutation bias shifts.

## INTRODUCTION

Mutation is the major source of genetic variation, and it is important to quantify the phenotypic and fitness effects of new mutations. A substantial body of work has therefore focused on determining the statistical distribution of mutational effects (the distribution of fitness effects, DFE) and the evolutionary processes that shape the DFE [1–3]. The DFE determines the number and proportion of beneficial mutations, a key parameter in population genetic models of evolutionary change. A broad and general understanding of the DFE and factors that influence it is thus crucial to predict adaptation rates, trajectories, and fates of evolving populations. Ultimately, such predictions and their tests are key to tackling problems such as the emergence of antimicrobial resistance and rapid environmental change that threatens populations [4] . From numerous studies using different approaches to estimate or quantify the DFE [5], we know that it is influenced by several factors such as the genetic background, the environment, the effective population size, and prior history of adaptation [1–3,6,7].

New work in the past few years has suggested that the DFE may also vary with the nature of the underlying mutations. The mutation spectrum – describing relative frequencies of different types of mutations – is typically biased towards specific classes of mutations. For instance, most organisms show a bias towards more transition mutations [8]. If different mutational classes have distinct fitness consequences, such pervasive mutation biases may affect the DFE. For example, an *Escherichia coli* strain that samples a higher proportion of AT→CG transversion mutations had a distinct DFE for antibiotic resistance, compared to a strain that samples more GC→TA transversions [9]. During laboratory evolution under increasing antibiotic stress, strains with different mutational bias developed resistance using distinct mutational paths [10]. Expanding this idea to genome-wide mutational biases, we recently showed that changing the mutation bias can alter the *E. coli* DFE across several environments with limiting carbon sources [11]. Specifically, on deleting a DNA repair gene (*mutY*) – which increases the incidence of transversion mutations compared to the wild type (WT) – we observed an ∼6% increase in the fraction of beneficial mutations on average across environments. The form of the global DFE may thus change depending on the mutation spectrum of an organism and the fitness effects of different types of mutations.

Simulations of adaptive walks as well as a mathematical model uncovered general conditions under which mutation bias shifts should change the DFE, and the evolutionary impacts of the expected DFE changes [11,12]. This led to a key prediction: opposing (i.e., reversing or reducing) the direction of the ancestral mutation bias should increase the fraction of beneficial mutations (Figure 1). Opposing an existing mutation bias is predicted to be generally beneficial because it allows populations to explore previously under-sampled mutational space, including beneficial mutations that were not fixed (and were therefore available). For instance, consider a population that has evolved in a constant environment with a transition mutation bias (e.g., WT *E. coli*) for some time. It will have gradually sampled, and fixed, many of the possible beneficial transition mutations; but it would have sampled only a small fraction of available beneficial transversions. Thus, adaptation results in a depletion of the beneficial part of the DFE (the DBFE), especially the well-sampled mutational classes [12–14]. On introducing a transversion bias (reversing the existing bias), such a population is more likely to sample beneficial transversions, leading to a right-shifted DFE (Figure 1). In contrast, if the ancestral bias is reinforced (i.e., with a stronger transition bias in *E. coli*), we predict that the DFE should shift left compared to the WT, with a larger fraction of deleterious mutations. Thus, mutation biases may play important roles in shaping DFEs [11,12] .

**Figure 1:**
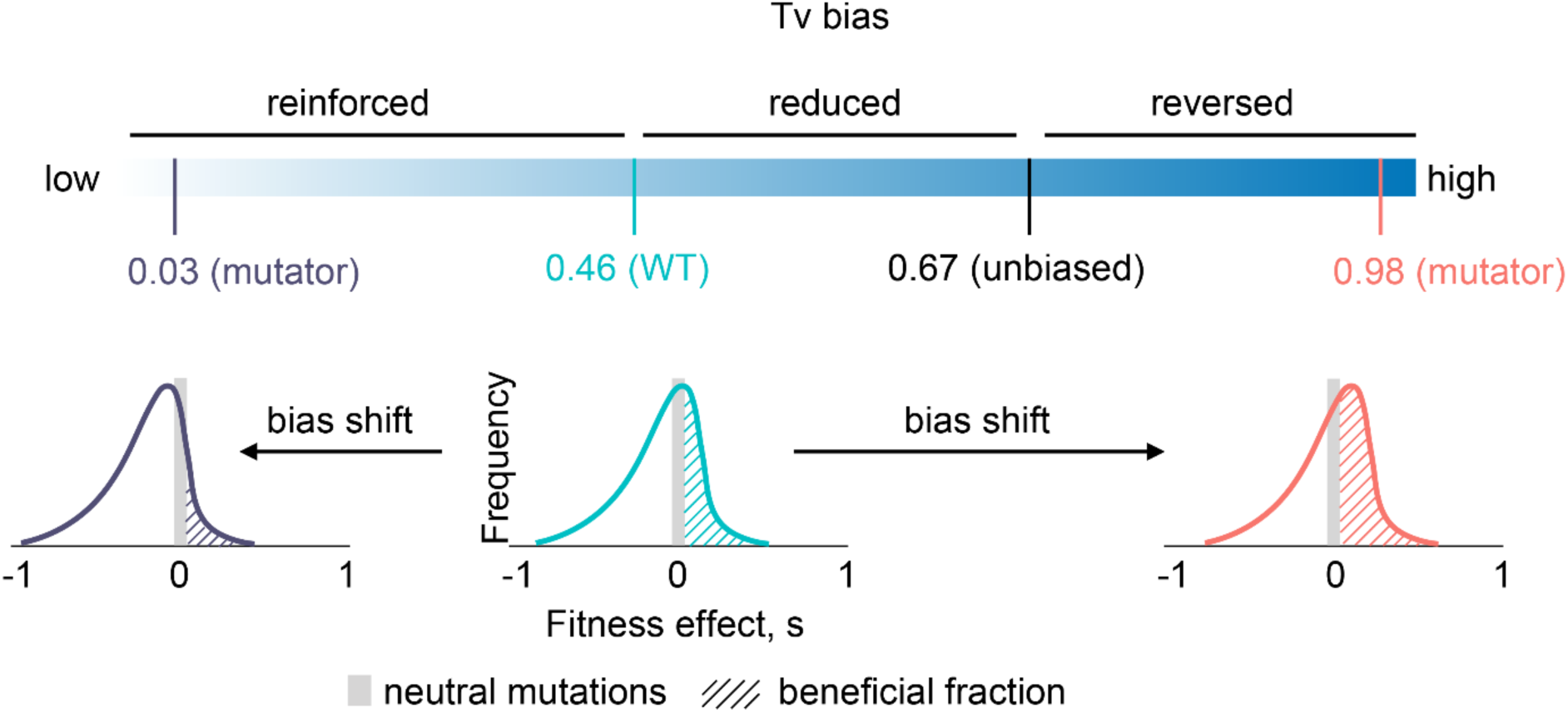
Schematic of the key prediction tested in this work. As the transversion (Tv) mutation bias of WT *E. coli* is shifted away from the ancestral bias, the resulting distribution of fitness effects (DFE) is predicted to change. Specifically, it should shift left with a bias reinforcement, with a lower fraction of beneficial mutations. In contrast, reversing the ancestral bias is predicted to cause right-shifted DFEs with higher proportions of beneficial mutations.

Here, we experimentally test these predictions by generating empirical DFEs of new mutations in six *E. coli* strains carrying deletions of various DNA repair genes involved in the mismatch repair (MMR) pathway, the 8-oxo-dGTP repair pathway, or the repair of damaged pyrimidines (Table 1). Wild type *E. coli* has a significant transition (Ts) bias, whereby only ∼46% of single nucleotide mutations are transversions (Tv) (compared to the null expectation of ∼67% Tv; Figure 1). Deleting DNA repair genes leads to a wide range of mutation biases, with transversion biases of 0.03 (i.e., 97% Ts) to 0.98 (i.e., 98% Tv) [8,15] . To obtain single-step mutations for constructing DFEs, we allowed several lineages of each strain to evolve independently in a mutation accumulation (MA) regime (some results from MA experiments for WT and Δ*mutY* were described in [11,16]). Evolution under MA allows nearly all mutations to be sampled (but see [17,18]), allowing us to obtain a random representative sample of mutations available to each strain. We used whole-genome sequencing to identify lineages with a single mutation compared to the respective ancestor, measured the fitness effect of each mutation in two environments (rich Lysogeny Broth (LB) and M9 minimal medium with Glucose), and used these data to construct DFEs. Our results provide strong support for our predictions, demonstrating that mutation spectrum shifts can determine the amount of adaptive genetic variation available to populations.

**Table 1:** Strains used in this study, and their mutational characteristics. For each class of mutations, bias is calculated as (no. of mutations of type A) / (no. of mutations of type A + no. of mutations of type B). Brief information about the function of deleted genes is provided in the footnotes [25] .

| Property | $\Delta\text{mutS}^\dagger$ | $\Delta\text{mutL}^\dagger$ | $\Delta\text{mutH}^\dagger$ | $\Delta\text{nth-nei}^\psi$ | WT | $\Delta\text{mutY}^\phi$ | $\Delta\text{mutT}^\phi$ |
| --- | --- | --- | --- | --- | --- | --- | --- |
| MA lines evolved | 350 | 350 | 350 | 300 | 98 | 430 | 300 |
| MA lines successfully sequenced | 345 | 344 | 346 | 285 | 97 | 424 | 271 |
| Total mutations in sequenced lines <sup>a</sup> | 616 | 603 | 908 | 502 | 426 | 798 | 789 |
| MA lines with single mutations <sup>b</sup> | 91 | 97 | 100 | 102 | 94 <sup>c</sup> | 113 | 97 |
| **Mutation rate** ($\times 10^{-8}$) | 1.4 | 1.4 | 2.1 | 0.17 | 0.0091 | 0.083 | 2.3 |
| $\pm 95\%$ CI | $\pm 0.14$ | $\pm 0.13$ | $\pm 0.14$ | $\pm 0.015$ | $\pm 0.0028$ | $\pm 0.016$ | $\pm 0.19$ |
| BPS | 508 | 509 | 809 | 482 | 387 | 787 | 783 |
| Indel | 108 | 94 | 99 | 20 | 39 | 11 | 6 |
| **Indel bias** | 0.18 | 0.16 | 0.11 | 0.04 | 0.09 | 0.01 | 0.01 |
| $\pm 95\%$ CI | $\pm 0.040$ | $\pm 0.038$ | $\pm 0.033$ | $\pm 0.023$ | $\pm 0.057$ | $\pm 0.011$ | $\pm 0.010$ |
| Coding | 440 | 437 | 710 | 400 | 282 | 684 | 657 |
| Noncoding | 68 | 72 | 99 | 82 | 105 | 103 | 126 |
| **Noncoding bias** | 0.13 | 0.14 | 0.12 | 0.17 | 0.27 | 0.13 | 0.16 |
| $\pm 95\%$ CI | $\pm 0.036$ | $\pm 0.037$ | $\pm 0.035$ | $\pm 0.044$ | $\pm 0.088$ | $\pm 0.032$ | $\pm 0.044$ |
| Non-synonymous | 290 | 274 | 433 | 273 | 183 | 489 | 462 |
| Synonymous | 150 | 163 | 277 | 127 | 99 | 195 | 195 |
| **Synonymous bias** | 0.34 | 0.37 | 0.39 | 0.32 | 0.35 | 0.29 | 0.30 |
| $\pm 95\%$ CI | $\pm 0.05$ | $\pm 0.051$ | $\pm 0.051$ | $\pm 0.054$ | $\pm 0.095$ | $\pm 0.043$ | $\pm 0.054$ |
| Transitions | 491 | 491 | 774 | 430 | 206 | 69 | 12 |
| Transversions | 17 | 18 | 35 | 52 | 178 | 644 | 771 |
| **Transversion bias** | 0.03 | 0.04 | 0.04 | 0.11 | 0.46 | 0.91 | 0.98 |
| $\pm 95\%$ CI | $\pm 0.019$ | $\pm 0.020$ | $\pm 0.021$ | $\pm 0.036$ | $\pm 0.099$ | $\pm 0.028$ | $\pm 0.015$ |
| AT→GC | 339 | 361 | 568 | 22 | 118 | 24 | 775 |
| GC→AT | 162 | 143 | 229 | 438 | 206 | 746 | 5 |
| No change | 7 | 5 | 12 | 22 | 60 | 17 | 3 |
| **GC→AT bias** | 0.32 | 0.28 | 0.29 | 0.95 | 0.64 | 0.97 | 0.01 |
| 95% CI | $\pm 0.049$ | $\pm 0.048$ | $\pm 0.048$ | $\pm 0.025$ | $\pm 0.096$ | $\pm 0.017$ | $\pm 0.01$ |

<sup>a</sup> Mutations used to calculate mutation rates and spectra.
<sup>b</sup> Lines used for fitness measurements and inference of DFEs.
<sup>c</sup> All of these single-step mutations were not derived from different MA lines (i.e., 80 of the 94 mutations were obtained from WT block 1 that contained only 38 lines, such that some lines contributed multiple single mutants). This is because 46 of them represent second-, third-, or fourth-step mutations occurring in a given lineage, whose fitness effect was estimated with respect to the immediate ancestor (i.e., carrying first-, second- or third- step mutations, respectively). Details are described in [16], and in Table S1.
<sup>†</sup> Genes involved in the methyl-directed mismatch repair (MMR) pathway. MutS recognizes mismatched base-pairs in double stranded DNA. MutL binds to DNA-bound MutS, and facilitates interaction with MutH. MutH performs daughter-strand recognition and creates single-strand DNA breaks in the daughter strand at the nearest hemi-methylated site. The breaks are then repaired by exonuclease activity extending up to the mismatch, followed by resynthesis by DNA polymerase III.
<sup>ψ</sup> Genes involved in the repair of damaged pyrimidines in double-stranded DNA. *nth* encodes Endonuclease III, which recognizes and cleaves N-glycosidic bonds of damaged pyrimidines, leaving abasic sites, and also cleaves the phosphodiester backbone 3' to the site to create single-strand DNA breaks. *nei* encodes Endonuclease XIII, and shares specificity with Endonuclease III.
- 1 $\Phi$ Genes involved in the 8-oxo-dGTP repair pathway. MutT hydrolyses cytosolic 8-oxo-dGTP to 8-oxo- - 2 dGMP, preventing its incorporation into DNA during replication. MutY recognizes 8-oxo-G:A mis-pairs - 3 in double-stranded DNA and excises the adenine, leaving an abasic site.

## METHODS

### Bacterial strains

We obtained the wild-type (WT) strain of *E. coli* K-12 MG1655 from the Coli Genetic Stock Centre (CGSC, Yale University), streaked it on Luria Bertani (LB) agar (Miller), and chose one colony at random as the WT ancestor for subsequent experiments. We similarly obtained the mutator strains of *E. coli* (Δ*mutT*, Δ*mutH*, Δ*mutL*, Δ*mutS*, Δ*nth,* Δ*nei*, Δ*mutY*) from the Keio collection of gene knockouts from CGSC (BW25113 strain background [19]). These gene knockouts were made by replacing open reading frames with a Kanamycin resistance cassette, such that removing the cassette generates an in-frame deletion of the gene. The design of gene deletion primers ensured that downstream genes were not disrupted due to polar effects. For each mutator strain, we moved the knockout locus from the BW25113 background into the MG1655 (our WT) background using P1 phage transduction [20]. We then removed the kanamycin resistance marker by transforming kanamycin-resistant transductants with pCP20, a plasmid carrying the flippase recombination gene and ampicillin resistance marker. We grew ampicillin resistant transformants at 42 °C in LB broth overnight to cure pCP20 and streaked out 10 µL of these cultures on LB plates. After 24 hours, we replica-plated several colonies on both LB + kanamycin agar plates and LB + ampicillin agar plates, to screen for the loss of both kanamycin and ampicillin resistance. We PCR-sequenced the knockout locus to confirm removal of the kanamycin cassette. For generating the Δ*nth-nei* double knockout, we first created a Δ*nth* strain, and then moved the Δ*nei* locus into this background using P1-phage transduction as described above. In the process of making gene knockouts, all mutator strains (except Δ*nth-nei*) acquired background mutations due to multiple generations of growth that occurred during the screening process. These are listed in Supplementary Datasheet S1.

### Mutation accumulation (MA) experiments

We used mutation accumulation (MA) experiments of varying length to obtain mutator strains carrying single mutations that would reflect the mutation spectrum of each mutator (∼100 per strain). We isolated a single colony of each ancestor, suspended it in LB broth, and plated it to obtain as many colonies as were needed for each independent MA line. We used this same broth culture within 3-4 hours of growth to extract DNA for whole genome sequencing (see next section). For mutators with very high mutation rates (ΔmutH, ΔmutT, ΔmutL and ΔmutS), each MA block had its own ancestor (since each time cells are grown up from freezer stocks, there is a very high probability that new mutations will arise) whose sequence was used to subtract the ancestral mutations from offspring lines (see below). For mutators with intermediate mutation rates and for WT, different MA blocks of a strain were started with the same ancestor.

The MA protocol minimises the effect of selection, allowing sampling of a wide range of mutational effects, largely independent of their fitness consequences. For each MA line, every 24 hours we streaked out a random colony (closest to a pre-marked spot) on a fresh LB agar plate. For MA experiments lasting more than a day, every 4-5 days we inoculated a part of the transferred colony into LB broth at 37°C for 2-3 hours and froze 1mL of the growing culture with an equal amount of 60% glycerol at –80°C. For 1-day MA experiments, we similarly cultured and froze the final chosen colony. We used these freezer stocks of the MA lines for sequencing. We chose the length and number of replicate lines of the MA experiments for each strain depending on mutation rate and logistical feasibility. Since the WT had a relatively low mutation rate, we founded a small number of MA lines to make daily transfers feasible, but evolved them for many generations. For the mutators, we founded larger numbers of MA lines but evolved them for just a few generations. We also split the large number of MA lines into blocks (except Δ*mutT*; Table S1) to make transfers and handling easier. Thus, most MA experiments were performed across at least two experimental blocks (Table S1).

We founded multiple MA lines of each strain from single colonies: WT (98 lines), Δ*mutT* (300 lines), Δ*mutH* (350 lines), Δ*mutL* (350 lines), Δ*mutS* (350 lines), Δ*mutY* (430 lines) *and* Δ*nth-nei* (300 lines) and propagated them through daily single-colony bottlenecks on LB agar plates (Table 1). We previously showed that our WT strain goes through ∼27 generations in 24 hours of growth on LB agar [16]. We used this estimate to calculate the number of generations elapsed in our MA experiments. We evolved WT lines in two experimental blocks: one block of 38 lines evolved for 300 days (8250 generations, described previously in [11,16]), and a second block of 60 lines evolved for 85 days (2295 generations) (Table S1). We evolved mutators with intermediate mutation rates in short MA experiments: Δ*nth-nei* in two blocks (block 1: 80 lines, 8 days, ∼216 generations; block 2: 220 lines, 8 days, ∼216 generations), and Δ*mutY* in three blocks (block 1: 300 lines, 12 days, ∼314 generations, described previously in [11] ; block 2: 80 lines, 5 days, ∼135 generations; block 3: 50 lines, 1 day, ∼27 generations; Table S1). Finally, we evolved mutators with very high mutation rates in very short MA experiments: Δ*mutS* (block 1: 300 lines, 1 day, ∼27 generations; block 2: 50 lines, 1 day, ∼27 generations), Δ*mutL* (block 1: 300 lines, 1 day, ∼27 generations; block 2: 50 lines, 1 day, ∼27 generations), Δ*mutH* (block 1: 300 lines, 1 day, ∼27 generations; block 2: 50 lines, 1 day, ∼27 generations) and Δ*mutT* (block 1: 300 lines, 1 day, ∼27 generations; Table S1).

### Whole-genome sequencing to identify clones with single mutations

We sequenced individual colonies from the MA experiments to identify all clones carrying a single mutation relative to their ancestor. For WT, Δ*nth-nei* and Δ*mutY*, we inoculated 2µL of the frozen stock of each evolved MA line into 2mL LB broth, and allowed the cells to grow overnight at 37°C with shaking at 200 rpm. For Δ*mutT*, Δ*mutH*, Δ*mutL* and Δ*mutS*, we allowed cells from frozen stocks to only grow for 3-4 hours, to minimize the accumulation of additional mutations. Next, we extracted genomic DNA (GenElute Bacterial Genomic DNA kit, Sigma-Aldrich) and quantified it using the Qubit HS dsDNA assay kit (Invitrogen). We prepared paired-end libraries from each line and the respective MA ancestors, and sequenced them on an Illumina platform (either 2×100 bp, or 2×125 bp, or 2×250 bp). Table S1 provides details of the library preparation and sequencing methods used, as well as the sequencing depth achieved for each strain. Whole genome sequencing (WGS) for some MA lines was unsuccessful, i.e., we obtained a very small number of reads or no reads at all; these lines were excluded from further analyses (Table 1, Table S1).

For each sample where WGS was successful, details of the number of mutations called are given in Supplementary Datasheet S2. We aligned reads with average quality score > Q30 to the NCBI reference *E. coli* K-12 MG1655 genome (RefSeq accession NC_000913.2) using the Burrows-Wheeler short-read alignment tool BWA [21], and used SAMtools to further process the BWA outputs and generate pileup files [21]. Next, we used the default parameters in the VarScan package [22] to extract lists of base-pair substitutions and short indels (<10-bp length). We used Breseq with default parameters [23] to identify long indels and duplications. From these mutation lists, we only retained mutations that satisfied the following 3 criteria: (i) mutations represented on both the plus and the minus strand, (ii) mutations supported by at least 4 reads per strand, and (iii) mutations with frequency > 80%. The first two filters would remove mutations with weak support, and the last filter would remove mutations that may have arisen either during the late stages of colony growth in MA experiments, or during the brief period of growth for stock preparation or DNA extraction. We performed this filtering using custom scripts written in R. Finally, we used a custom R script to remove mutations present in the corresponding ancestor from the mutation list of each evolved line, and generated the final mutation list for each lineage (Supplementary Datasheet S3).

We used several measures to maximize and estimate the accuracy and reliability of mutation-calling. To identify ancestral mutations, we sequenced ancestral MA clones at higher depth (Table S1), and relaxed the 80% frequency filter such that we captured ancestral mutations segregating at lower frequency. This was especially important for strains with very high mutation rate. We confirmed that all offspring MA lines seeded by a given ancestral clone showed the expected set of ancestral mutations at very high frequency. To quantify the potential effects of secondary low-frequency mutations on our fitness measurements (and therefore the DFE), for each evolved MA line we generated a separate list of mutations after relaxing the 80% frequency filter. We used the number and frequency of these secondary mutations to estimate the robustness of our results (described in the Results section). To determine the false negative rate of our WGS pipeline, we measured the recall rate of two known mutations in our WT ancestor (the progenitor of all our mutators) relative to the NCBI reference genome (RefSeq accession NC_000913.2) [16]: a G → A SNP at position 2845011 and a 2-bp CG insertion at position 4296380, expecting these mutations to be called at 100% frequency in all evolved MA lines. Finally, to determine the false positive rate, we measured the fitness of a subset of single-mutation clones of WT across two growth cycles, expecting a strong positive correlation if the identified mutations were real.

### Estimating mutation rate, spectra, beneficial supply, and deleterious load

We estimated the mutation rate (μ) and mutation bias for each mutator and WT using mutations called from all sequenced isolates (Table 1, row “MA lines successfully sequenced”). We calculated μ (per bp per generation) as:

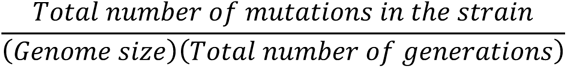

The number of mutations are given in Table 1 and the genome size is 4.6 x 10^-6^. The number of generations was calculated as:

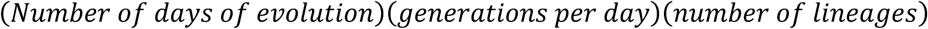

We tested the observed frequency distribution of the number of mutations called in each lineage, against the expected Poisson distribution for random mutations. We calculated mutational biases from the different types of mutations observed in the MA-evolved lines (Table 1) as described in [11]. For instance, we calculated the Tv bias as:

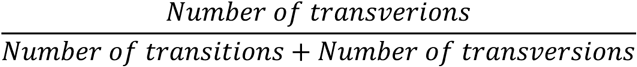

We estimated 95% confidence intervals for these estimates as 1.96 times the standard deviation of calculated bias. We estimated the confidence intervals for mutation rate estimates as the known mean mutation rate ± the margin of error for a t-distribution with known mean and unknown standard deviation.

For each strain, we used the estimated mutation rates and the fractions of beneficial and deleterious mutations (f_b_ and f_d_, see below) to calculate the predicted total supply of beneficial mutations as:

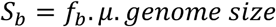

and the total genetic load due to deleterious mutations as:

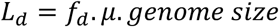

We estimated S_b_ and L_d_ using either the WT DFE (i.e., assuming that DFEs were invariant across strains), or using strain-specific DFEs measured in each environment. In each case we also estimated confidence intervals as 1.96 times the standard deviation of S_b_ or L_d_. We then compared these values for each strain, to quantify the impact of mutation bias on the supply of beneficial mutations and the deleterious load.

### Growth rate measurements to construct single-mutation DFEs

From the complete set of all MA-evolved lines with WGS, we focused on those that had a single new mutation compared to the respective ancestor. This included 91 clones of Δ*mutS*, 97 Δ*mutL*, 100 Δ*mutH*, 102 Δ*nth*-*nei*, 94 WT (80 from block 1 [16] and 14 from block 2), 113 Δ*mutY* (79 from block 1 [11], 26 from block 2 and 8 from block 3), and 97 Δ*mutT* MA lines (Table 1, row “MA lines with single mutations”). We performed all fitness assays and subsequent analyses (described below) with these 694 isolates. We measured growth rates of each evolved MA line with a single mutation and its respective ancestor in two liquid culture media: LB broth (Miller, Difco) or M9 minimal salts (Difco) + 5 mM glucose. We inoculated each isolate from its freezer stock into either LB broth or M9 minimal salts medium with 5mM glucose, and allowed it to grow at 37 °C with shaking at 200 rpm for 14-16 hours. We inoculated 2 µL of this culture into 200 µL growth media in 96-well plates (Costar) and incubated the well plate in a Tecan F200 Multimode plate reader at 37°C with orbital shaking at 185 rpm for 16-18 hours. Every 15 minutes, the plate reader measured the optical density (OD600) for all wells. In each plate, we included the MA ancestors relevant for the evolved MA isolates to enable fitness calculations, a reference strain (the parent WT strain) to enable checks for consistency across plate reader runs, and blank control wells to check for media sterility. For each evolved isolate, we used the average growth rate of three technical replicates to calculate the relative growth rate as:

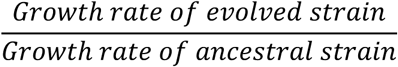

For WT MA lines, we used the WT ancestor; and for mutator MA lines, we used the mutator ancestor. We estimated maximum growth rate, obtained from a linear fit to log OD600 vs. time curves, using the Curve Fitter software [24] . The fitness effect of each mutation (*s*) was then calculated as:

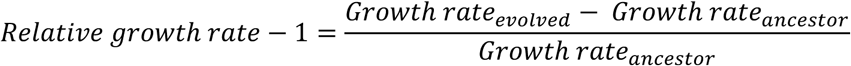

We used *s* values of mutations to construct strain- and environment-specific distributions of fitness effects (DFE). Importantly, we corrected our DFEs for the expected selection bias in bacterial colonies in MA experiments as described before [11,17] . Briefly, the correction involves binning the measured selection coefficients into discrete bins, and then down-weighting the beneficial fraction of the DFE and over-weighting the deleterious fraction to account for the slightly higher probability of finding beneficial mutations. The bias-corrected vs. uncorrected DFEs are shown in Figures S8 and S9; note that the bias correction procedure leads to discretized bins in the corrected frequency distributions (DFEs).

## RESULTS

### Accuracy of mutation calling and fitness measurements

As described above, our main goal in this study was to construct and compare single mutation DFEs across strains with distinct mutation biases. To maximize the accuracy of mutation calling, we used stringent sequencing quality filters (see methods), e.g., only using lineages with high sequencing depth (mean ∼40x across all lines, Table S1). We also conducted several analyses to test the accuracy and reliability of our WGS pipeline. We first confirmed that two known mutations in our WT ancestor (the parent of all our strains) were called in every single evolved MA line with high frequency (SNP at ∼100% allele frequency, indel with >93% read support; Figure S1), indicating a very low false negative rate. Second, in all strains, the observed frequency distribution of the number of mutations called in each lineage was indistinguishable from the expected Poisson distribution for random mutations (Figure S2), suggesting that non-random processes or mutation calling protocols did not significantly influence the outcome of our MA experiments. Third, we identified one-mutation clones only when they had a single mutation at >80% frequency; on relaxing this filter we found either no secondary mutations or secondary mutations segregating at low frequencies (Figure S3). Together, these results suggest high accuracy of mutation-calling in our study.

Next, we tested the repeatability and reliability of our fitness estimates, which entailed measuring exponential growth rates of each of the single-mutation clones. In previous work, we had found high repeatability of growth rates [11,16]. We re-confirmed this for the current study: for a subset of isolates, relative fitness measured by two different experimenters in different years was strongly positively correlated (Figure S4), as were fitness estimates in 48-well vs. 96-well plates performed in two different years (Figure S5). These analyses gave us confidence that our measured fitness values are generally robust. Finally, for single-mutation clones of WT, fitness values across two successive growth cycles were also strongly positively correlated (Figure S6). Together, these results indicate that the single mutations that we called were “real”, and that our fitness measurements were reliable.

### Transversion-biased strains have a right-shifted distribution of fitness effects (DFE) with a higher proportion of beneficial mutations and lower deleterious load

To test our hypothesis that transversion-biased *E. coli* strains can access more beneficial mutations, we constructed single-mutation DFEs for strains with different transition/transversion biases (Table 1). For each strain, we characterized the fitness effects of a total of ∼100 single mutations in two growth media: one rich (LB) and one relatively poor (minimal medium with glucose), using exponential growth rate as a measure of fitness. Using these fitness estimates, we first constructed raw DFEs, and then corrected them to account for selection bias during MA (Figures S7, S8). Note that the bias correction does not alter the selection coefficients measured for each mutation, but directly modifies the DFE to down-weight the proportion of beneficial mutations (see Methods). The median selection coefficient of single mutations (estimated from the corrected DFEs) varied from –0.175 to +0.025 (i.e., a 17.5% reduction to 2.5% increase in growth rate) in LB, and from –0.013 to +0.063 (i.e., 1.3% reduction to 6.3% increase in growth rate) in glucose.

As predicted, in both environments, strains that reinforced the WT (ancestral) mutation bias (i.e., were strongly Ts-biased: Δ*mutS*, Δ*mutL*, Δ*mutH* and Δ*nth-nei*) had left-shifted DFEs relative to WT (Figures 2 and 3). In contrast, the DFEs of strains that opposed the WT mutation bias (i.e., had a strong Tv bias, Δ*mutY* and Δ*mutT*) had relatively right-shifted DFEs. The DFE differences across strains were reflected in the proportion of beneficial mutations (f_b_), which was significantly higher in Tv-biased vs. Ts-biased strains (Figure 4A; Table S2, Table S3). Concomitantly, the fraction of deleterious mutations (f_d_) was significantly lower in Tv-biased strains (Figure 4A; Table S2, Table S3). These patterns did not change when we imposed more stringent conditions for single-mutation calling, e.g., if we constructed DFEs using only clones with no secondary mutations or with low-frequency secondary mutations, or clones with two mutations (Figure S9). Similarly, reducing sample sizes commensurate with the stringent filtering also did not alter the patterns (Figure S10). Thus, the results are robust, and support our main prediction (Figure 1) that mutation bias shifts that reinforce the Ts bias of WT *E. coli* will have left-shifted DFEs and those that reverse the bias (i.e., are transversion biased) will have right-shifted DFEs (Figure 4A). A related prediction is that the magnitude of the DFE shift is correlated with the magnitude of the bias shift, and this is also consistent with our results (Figures S11A–B, f_b_ increases significantly with increasing Tv bias; Figure S11C– D, f_d_ tends to reduce with Tv bias, but not significantly so). However, we caution that given the limited number of bias-shifted strains in our dataset, additional work is required to adequately test this correlation. Despite the overall trends described above, we note some exceptions. In LB, Δ*mutY* had a much higher f_b_ compared to Δ*mutT*, despite a slightly weaker Tv bias (Figures 4A–B; Table S2). In glucose, Δ*mutL* and Δ*mutH* had higher f_b_ values despite the same transition bias as Δ*mutS* and a stronger Ts bias than Δ*nth*-*nei* (Table S3). We explore these exceptions in more detail in the discussion section.

**Figure 2:**
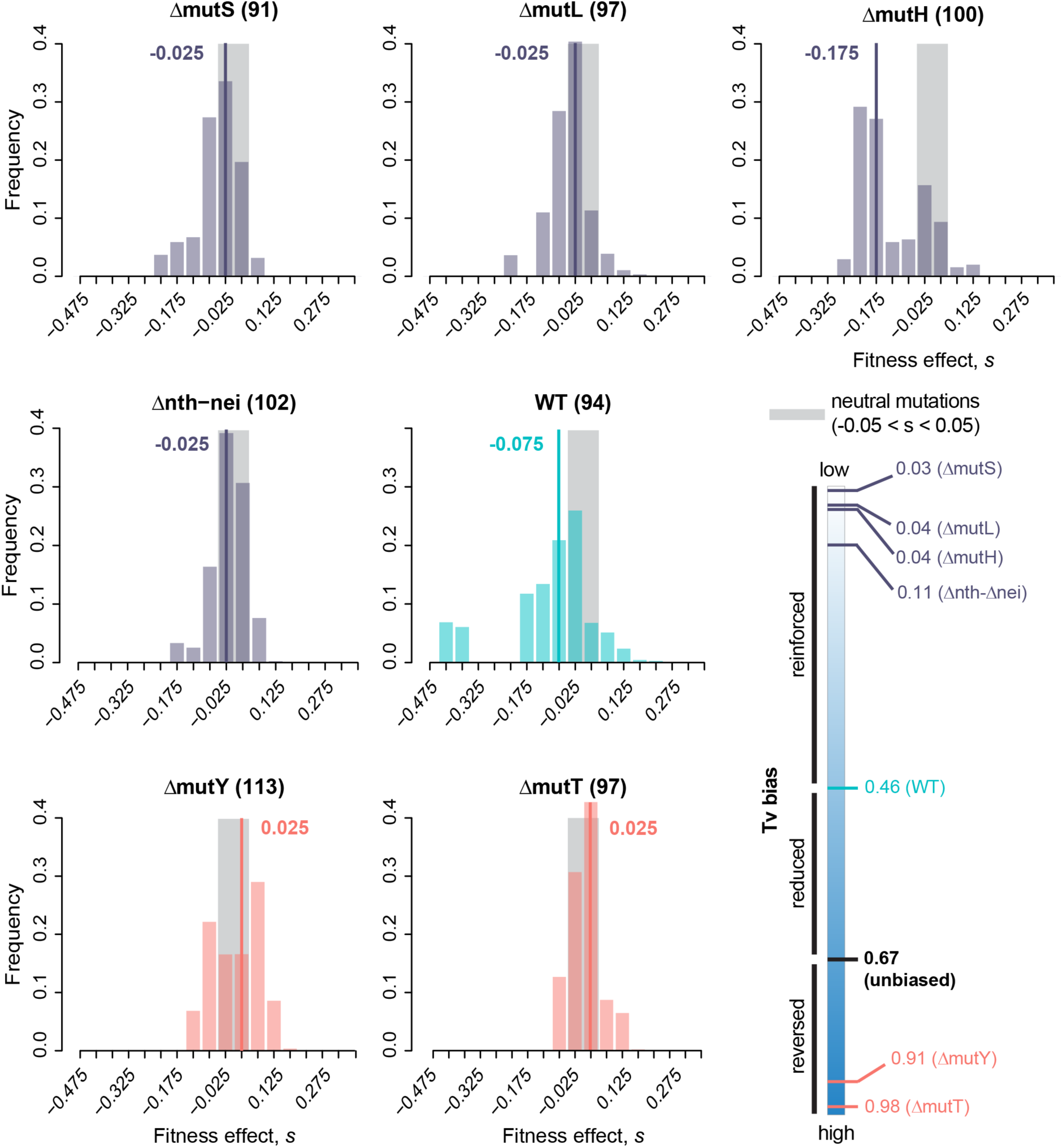
Distribution of fitness effects of new mutations in a rich medium (Luria Bertani broth, LB) for *E. coli* strains with varying mutation bias. The distribution of fitness effects of single randomly occurring mutations in each strain (sample size in parentheses), calculated as maximum growth rate relative to the respective ancestor (x-axis). The schematic at bottom right shows the transversion (Tv) bias of each strain. The WT strain is shown in cyan; strains that reinforce the WT transition bias are in purple; strains that reduce or reverse the WT bias (i.e., have a transversion bias) are in pink. Each DFE was corrected for selection bias; raw (uncorrected) DFEs are shown in Figure S7; raw fitness values are provided in Supplementary Datasheet S3. Grey areas indicate neutral mutations (*s*=0±0.05 to account for experimental measurement error); bold lines and numbers indicate median values of *s*.

**Figure 3:**
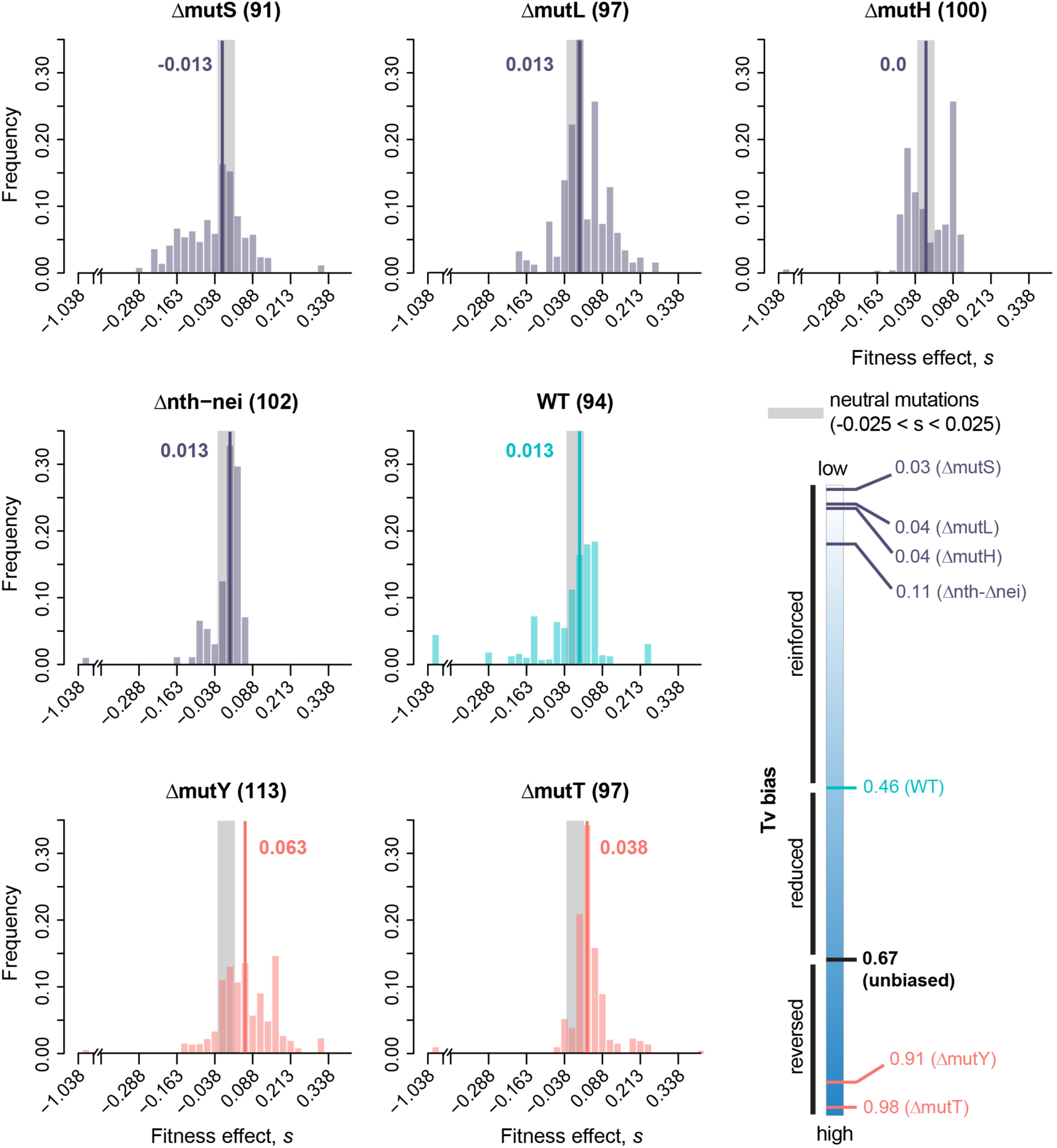
Distribution of fitness effects of new mutations in a minimal medium (M9 broth with 5 mM glucose) for *E. coli* strains with varying mutation bias. The distribution of fitness effects of single randomly occurring mutations in each strain (sample size in parentheses), calculated as maximum growth rate relative to the respective ancestor (x-axis). The schematic at bottom right is identical to the schematic in Figure 2, and shows the transversion (Tv) bias of each strain. The WT strain is shown in cyan; strains that reinforce the WT strain’s transition bias are in purple; strains that reduce or reverse the WT bias (i.e., have a transversion bias) are in pink. Each DFE was corrected for selection bias during MAClick or tap here to enter text.; raw (uncorrected) DFEs are shown in Figure S8; raw fitness values are provided in Supplementary Datasheet file S3. Grey areas indicate neutral mutations (*s*=0±0.025 to account for experimental measurement error); bold lines and numbers indicate median values of *s*.

**Figure 4:**
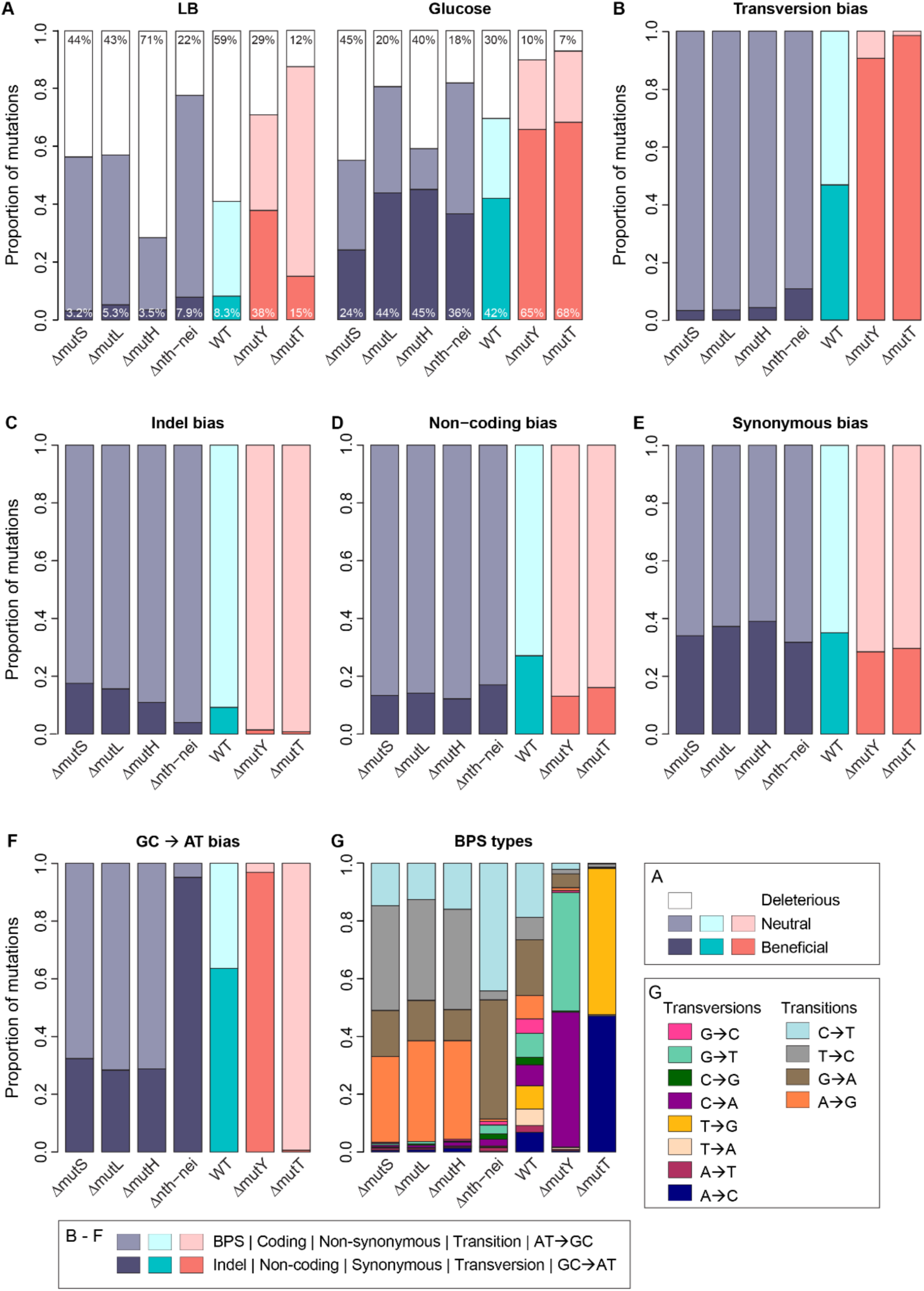
Differences in DFEs of strains are most strongly associated with transversion bias. In panels A—F, bars are colored by the mutation bias of each strain, as indicated in Figures 2 and 3. (A) Stacked bar plots show the total fraction of neutral, deleterious, and beneficial mutations observed in the DFEs of each strain in each growth medium, extracted from the DFEs shown in Figures 2 and 3. Percentage deleterious and beneficial mutations are indicated at the top and bottom of each bar respectively. Panels B–G show different aspects of the mutation spectra of strains, with darker vs. lighter shades indicating mutational classes. (B) Tv bias (C) Indel bias (D) Noncoding mutation bias (E) Synonymous mutation bias (F) GC→AT bias (note that mutations that do not affect GC→AT bias, i.e., AT→TA and GC→CG mutations, are not shown here) and (G) Types of base-pair substitutions (BPS).

An important consequence of increased genomic mutation rates is that mutators enjoy an increased total beneficial mutation supply (S_b_), but must also contend with higher deleterious genetic load (L_d_) – both important factors in determining their fates during adaptation [26–28]. We first calculated S_b_ and L_d_ for all strains assuming WT f_b_ and f_d_ (i.e., assuming similar DFEs across strains) [29]. As expected, these values scale linearly with the genomic mutation rate, with strains with 100x higher mutation rates predicted to have a 10–250-fold greater beneficial supply and lower deleterious load compared to WT (Figure S12, Tables S4–S5). However, on accounting for the altered f_b_ and f_d_ values of mutator strains due to their DFE shifts, we observed deviations from this relationship. Ts biased strains typically had lower S_b_ while Tv biased strains had higher S_b_ than expected (Figure S12A and S12B, compare open circles vs. filled circles; Table S4). In the case of L_d_, accounting for the observed DFEs caused relatively small changes for Ts biased strains, whereas Tv biased strains had a substantially lower L_d_ than expected based on mutation rate alone (Figures S12C and S12D, Table S5). The effect of using the empirically observed DFEs was strongest for Δ*mutT*, where the beneficial supply relative to wild type increased from ∼250 fold to ∼650 and ∼400 fold in LB and glucose respectively (Table S4), and the deleterious load reduced from ∼250 fold to ∼53 fold in LB and ∼58 fold in glucose (Table S5). Thus, mutation bias shifts could have very large effects on the evolutionary fate of mutators.

### DFE shifts and fraction of beneficial mutations are strongly associated with reversal of Tv bias, with some unexplained variation

The strains used in our analysis differed in several aspects of their mutation spectra (Table 1), so we examined variation in each aspect of the spectrum in more detail (Figures 4B–G). The increase in f_b_ across strains was positively correlated with the magnitude of Tv bias (i.e., stronger bias reversal) (Figures S11A–B; also compare Figures 4A and 4B). In contrast, no other axis of variation in mutation spectrum matched the pattern of variation in f_b_ across strains (compare Figure 4A with Figures 4C–F). In each of these cases, either the range of variation in mutation bias across strains was very small (indel bias, synonymous bias, non-coding bias; Figures 4C–E, Table 1) or there was no consistent pattern (noncoding bias and GC→AT bias; Figures 4D and 4F respectively). Even when we considered each type of base substitution separately, the fitness effects and type of mutation were not associated (Figure 4G). For instance, Δ*mutY* and Δ*mutT* each sample distinct types of transversion mutations, yet both have the highest f_b_ values in both environments. Finally, pooling data across all strains in our dataset, transversion mutations were significantly more beneficial than transitions (Figures 5A and 5B), whereas no such fitness difference was observed for any other aspect of the mutation spectrum (except BPS vs. indels in glucose; Figure S13). Together, these results pointed to transversion bias as the dominant cause of DFE differences across strains.

**Figure 5:**
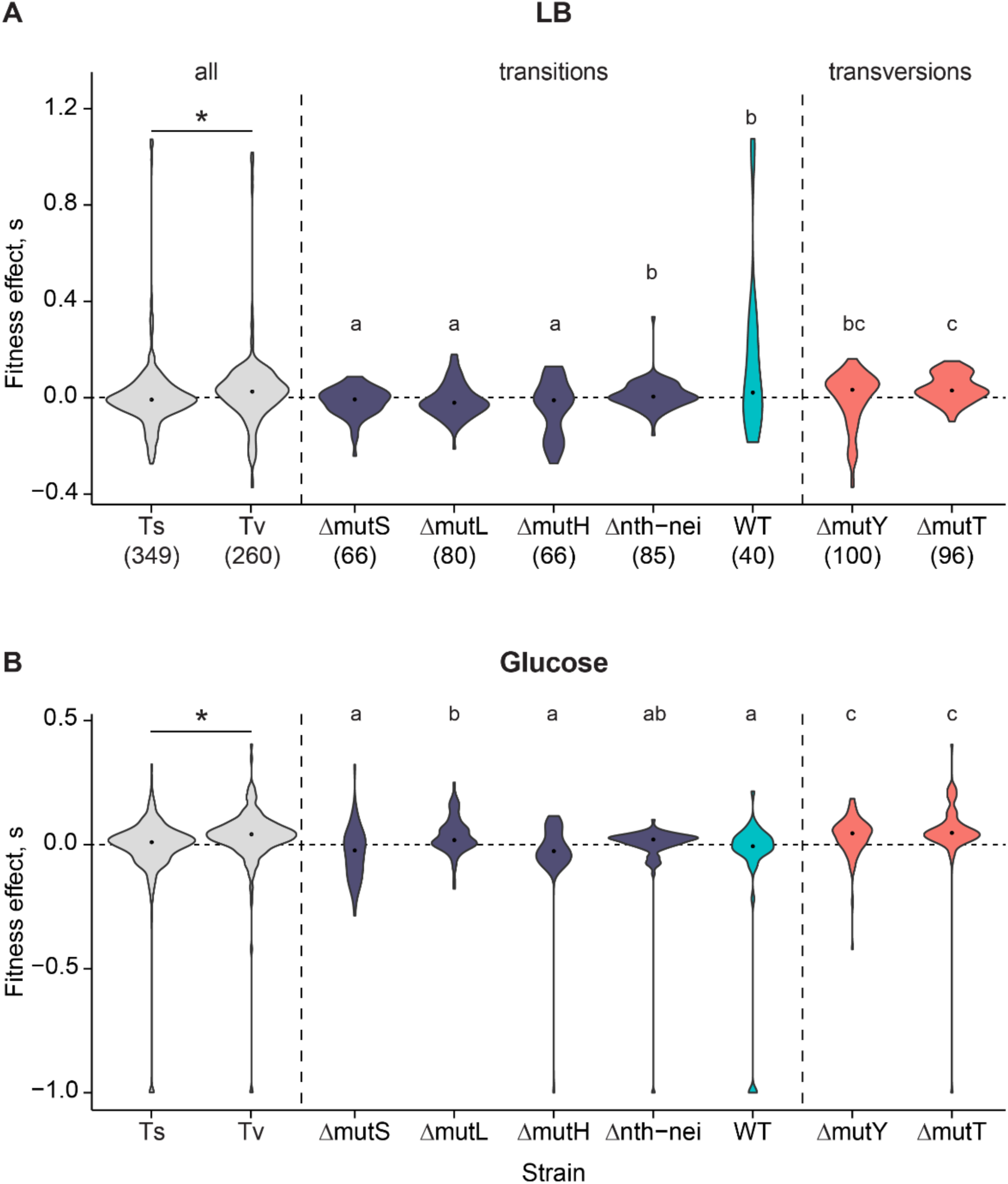
Transversion mutations are more beneficial than transition mutations. Fitness effects of mutations in (A) LB and (B) glucose; sample sizes are shown in x-axis labels in panel A. Note that y-axis ranges differ across panels. In each panel, the first two boxplots compare the effects of all transition vs. transversion mutations, pooled across all strains (asterisks indicate significant differences in Wilcoxon’s rank-sum tests; LB: p = 7.7E-10; Glucose: p = 5.1E-10). Following this, boxplots show the fitness effects of only Ts mutations from transition-biased strains and only Tv mutations from transversion-biased strains. Strains with the same letter have similar fitness effects based on pairwise Wilcoxon’s rank sum tests with Benjamini-Hochberg correction (i.e., all strains marked “a” have similar fitness, and all are significantly different from strains marked with “b” or “c”. Strains marked “bc” are similar to both “b” and “c”). Comparisons across all other types of mutations are shown in Figure S13.

Next, we tested whether the observed associations between transition/transversion bias and fitness effects of mutations are confounded by strain background (i.e., specifically which DNA repair genes were deleted and what other mutations occurred during the genetic manipulations (Supplementary Datasheet S1). We first tested whether higher initial fitness of each strain was associated with lower f_b_, reflecting the expected pattern of diminishing beneficial mutations with increasing background fitness [30,31] . Although the fitness of the original deletion strains varied significantly in both media, it was not correlated with f_b_ (Figure S14). We then conducted pairwise comparisons between strains, considering only Ts and Tv mutations. We expected that Tv mutations should be generally more beneficial than Ts mutations regardless of strain background, and the same type of mutation (Ts or Tv) should have similar fitness effects in all strain backgrounds. These predictions are broadly borne out in both media (Figures 5A and 5B): Tv mutations in Δ*mutY* and Δ*mutT* had similar (and higher) *s* values than Ts mutations, though in LB the fitness effects of Tv in Δ*mutY* were similar to Ts in WT and in Δ*nth*-*nei*.

Notably, in some cases Ts mutations in different Ts-biased strains had significantly different fitness effects (Figures 5A and 5B). In LB, Ts mutations in WT and Δ*nth-nei* strains were more beneficial than other Ts-biased strains, and in glucose, Ts in Δ*nth*-*nei* and Δ*mutL* were more beneficial. These exceptions could potentially be explained by aspects of the mutation spectra other than Ts/Tv bias, but these strains do not stand out as exceptional along other axes of the mutation spectrum (Figures 4B–G, Table 1). Further, comparing Ts-biased strains in each growth medium, we found very few and inconsistent differences in fitness effects along other axes of mutation bias (e.g., coding vs. non-coding, synonymous vs. non-synonymous; Table S6), indicating that they cannot explain the differences in fitness effects across Ts-biased strains. To summarize, the broad patterns of DFE variation that we observed are explained by Ts/Tv bias, but there is additional variation among Ts-biased strains that remains unexplained.

### Reversal of the GC/AT bias does not alter DFEs

Prior simulations had predicted that the beneficial effects of sampling unexplored mutational space should extend to any axis of the mutation spectrum, including the GC→AT bias [11] . However, here we did not observe the predicted impacts of reversing the GC→AT bias. The WT has a slight GC→AT bias relative to the unbiased expectation of 0.5, and all three MMR strains as well as Δ*mutT* reverse this bias, whereas Δ*mutY* and Δ*nth-nei* strongly reinforce the bias (Table 1, Figure 4F). However, as discussed above, the DFEs of the strains were not correlated with the magnitude of GC→AT bias, and overall, the fitness effects of GC→AT mutations vs. AT→GC mutations were not distinguishable (Figures 6A–B). Since the GC→AT bias varies substantially across genes (Figures S15A–B), we hypothesized that local rather than global bias reversal may be more relevant. However, mutational effects were not correlated with gene GC content regardless of the direction of mutation bias (Figures 6C–D; also see Figures S15C–F). Thus, in contrast to the effects of reversing the Tv bias and contrary to our previous simulations, neither local nor global GC bias reversal altered the fitness effects of new mutations.

**Figure 6.**
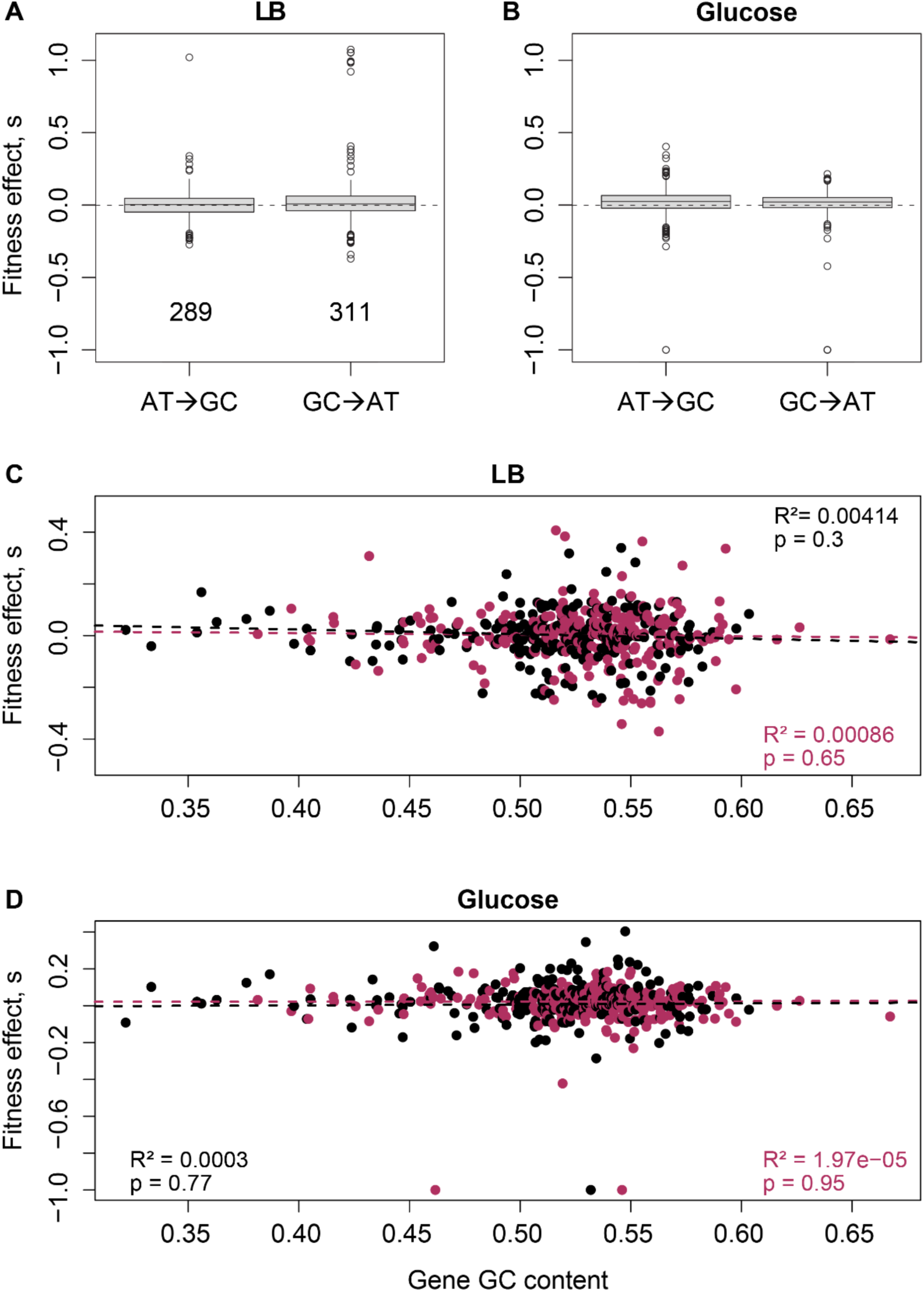
Reversal of the GC/AT bias does not impact fitness effects of mutations. Boxplots show fitness effects of AT→GC vs. GC→AT mutations in (A) LB and (B) Glucose. In each plot, data are pooled across all strains; sample sizes (total number of single mutations tested) are shown in the LB panel. 9 mutations that did not alter GC content (i.e., GC→CG or AT→TA mutations) are not shown here. Mutation type did not change fitness effects in either panel (Wilcoxon’s rank-sum test; LB: p = 0.4; Glucose: p = 0.24. (C–D) Scatter plots show the relationship between gene GC content and fitness effects of either AT→GC mutations (maroon points) and GC→AT mutations (black points) in (C) LB and (D) Glucose.

## DISCUSSION

Our results represent the first systematic experimental analysis of the fitness consequences of varying mutation bias, and support the prediction [11,12] that a reversal of an ancestral mutation bias (here, Ts bias in *E. coli*) can lead to a large increase in beneficial mutations. The difference is stark, with Tv-biased strains showing ∼2.5 to 12 times higher f_b_ relative to Ts-biased strains (Figure 4A). Note that even a small difference in f_b_ may alter the dynamics of adaptation in large asexual populations, by increasing the overall beneficial mutation supply (e.g., [32]). These results substantially expand upon our prior work, where we observed significantly higher f_b_ in Δ*mutY* compared to WT in 9 of 16 carbon environments [11]. One unexplained result in the previous study was the lack of significant differences in f_b_ in five environments, including LB and minimal glucose media. However, in our current analysis with increased sample sizes (80 vs. 94 for WT and 79 vs. 113 for Δ*mutY*), we observed significantly higher f_b_ in both LB and glucose, suggesting that the previously observed inconsistency across environments may have resulted from low statistical power. Another difference between the two studies is that fitness was measured in slightly different experimental contexts, in 48-well (older study) vs. 96-well microplates (current study). However, this cannot explain the different outcomes for WT and Δ*mutY*, because fitness is strongly positively correlated across microplate types (Figure S5). Thus, we suggest that with sufficiently large sample sizes, a Tv bias should consistently lead to right-shifted DFEs in diverse environments for historically Ts-biased species such as *E. coli*.

More generally, our results add to the growing realization that mutation bias can significantly shape evolutionary dynamics [33–35], by providing direct experimental evidence and suggesting a mechanism through which specific types of bias shifts may alter evolutionary outcomes (i.e., bias reversals lead to right-shifted DFEs with more beneficial mutations). Earlier work had also suggested that bias shifts can dramatically alter evolutionary dynamics, but the underlying mechanism and interpretations were different [9,10] . These studies showed (using some of the same strains used in our work) that distinct mutation spectra endow strains with differential success in sampling specific antibiotic resistance mutations. As a result, some strains are predicted to adapt faster to specific antibiotics. However, this explanation is specific to particular antibiotics and the relevant mutational targets of resistance. Thus, predicting the relative success of different strains in a given antibiotic would require knowledge of resistance mechanisms and whether they are more likely to arise via a specific type of mutation. In contrast, we provide a more general explanation, predicting that the effect of mutation bias depends on the prior evolutionary history of the population, such that a large shift favoring a previously poorly sampled class of mutations should be generally advantageous [11,12] . We hope that future work will test the effect of specific matches between selection and mutation bias vs. a broad bias reversal, independent of the source of selection. Such analyses are critical to expand our ability to predict adaptive outcomes and mutation fates under diverse selection pressures.

Although our experiments support our prediction about the impact of mutation bias reversals on the DFE, there are some interesting points of divergence. For instance, in LB, Δ*mutY* has a larger-than-expected f_b_, and the reason is not yet clear. Most intriguing is the variation in the DFEs of three strains with near-identical Ts bias in glucose (Δ*mutH*, Δ*mutL*, Δ*mutS*; Figure 4A), where the same DNA repair pathway (mismatch repair, MMR) is disrupted. Clearly, mutation bias shifts alone cannot explain these differences. One reason could be that some of the MMR genes have other functions apart from DNA repair, such that their deletion directly influence the fitness effects of new mutations. Alternatively, despite attempts to minimize structural disruptions, some repair gene deletions may have caused regulatory changes in downstream genes, leading to strain-specific epistasis with new mutations. Although we did not observe such strain-specific epistasis for a set of 19 mutations placed in both WT and Δ*mutY* backgrounds [11], further experiments are necessary to test this hypothesis for genes in the MMR pathway, whose deletion leads to distinct DFEs. Another interesting exception to the effect of Tv bias reversal is that while the overall pattern of change in f_b_ and f_d_ is consistent in both media (LB and glucose), we do see media–specific effects. In previous work with WT and Δ*mutY*, we had also observed significant differences in DFEs across environments with different carbon sources [11]. While variation in DFEs across environments is not surprising, the mechanisms underlying change in the rank order of f_b_ value across environments remain unclear. Analyzing the causes of the differences in DFEs across strains with similar mutation bias, as well as across environments, is thus a fruitful avenue for further work.

Another unexplained pattern in our current study is that despite strong reversals and reinforcements of the wild type GC→AT bias in some strains, GC→AT vs. AT→GC mutations had similar fitness effects, and this aspect of the mutation spectrum did not explain variation in the DFE. This is puzzling because simulations had predicted that reversal of *any* aspect of the mutation spectrum should influence the DFE, specifically shown for GC→AT bias [11]. One potential explanation is that the local GC bias rather than genome-wide GC bias is more relevant (e.g., due to the distinct evolutionary history of different genes). A second related hypothesis is that the fitness effects of GC/AT mutations depend strongly on their impact on gene function (e.g., distinct protein-disruptive effects of GC→AT vs. AT→GC mutations), and these effects depend on the genome GC content [36] . However, since we do not see any fitness differences between GC→AT or AT→GC mutations globally or locally (in GC-rich or GC-poor genes), neither hypothesis is supported by our data. A third possibility is that the GC→AT bias varies more frequently and/or dramatically across evolutionary time, reducing the impact of experimentally introduced GC/AT bias reversals relative to Ts/Tv bias reversals. Prior work shows that GC bias is indeed quite dynamic in the bacterial phylogeny (e.g., [37]), but it is impossible to directly test whether GC bias shifts occur more frequently than Tv bias shifts, because the latter do not leave a genomic signature. However, it is possible to approximately estimate the frequency and direction of bias shifts using the gain and loss of DNA repair genes with known effects on Ts/Tv and GC/AT bias. Using such analysis, we had previously observed that reversals of GC/AT bias occurred more frequently than Ts/Tv in the bacterial phylogeny (see Fig. 5C in [11]). Thus, we speculate that more frequent changes in the genomic GC content may explain why GC bias reversals did not impact the DFE, and emphasize the need for further theoretical and empirical work on the relative effects of bias shifts along different axes of the mutation spectrum.

A striking result from our study is the high proportion of beneficial mutations observed in all strains in glucose, and in transversion-biased strains in LB (Figure 4A), consistent with our previous analysis of WT and Δ*mutY* strains during growth in several carbon sources [11]. As discussed previously by us and by others, such high f_b_ values are not as rare as generally believed, especially during mutation accumulation studies [11,38] . In a recent review, Bao and colleagues suggest that the widely variable outcomes of fitness decline in MA lines (an indicator of the proportion of deleterious vs. beneficial mutations) is not easily explained by organism, genetic background, or test environment, but may arise from complex interactions between these and/or other factors [38] . In our current study, we ruled out several mechanisms that could artificially inflate the observed f_b_ values: selection bias during MA (we corrected our DFEs for such bias), low ancestral fitness leading to high f_b_ values [30,31,39] (we did not find a correlation between ancestral fitness and f_b_), and the impact of specific strain backgrounds (we observe a general effect of Ts vs. Tv mutations on f_b_). We hope that further analyses will clarify this issue.

Our results also inform several other open questions in the field, such as the fitness consequences of different types of mutational classes. Perhaps most interesting is the lack of significant differences in the fitness effects of synonymous and nonsynonymous mutations (Figure S13C), adding to the already substantial body of work suggesting that this distinction is not as large or widespread as expected [40,41] . Most prior studies compared the impact of synonymous and non-synonymous changes in specific genes of interest, so our results complement these studies in showing similar fitness effects across hundreds of mutations across the genome. Similarly, we do not find significant differences in coding vs. noncoding mutations (Figure S13B), or AT→GC vs. GC→AT mutations (Figure 6A-B). Together, our results indicate that while the fitness consequences of each of these types of mutations is probably highly context-specific (i.e., within genes, the fitness effect of mutations varies across sites), such patterns are not observed at the genome-wide scale. Future studies with larger sample sizes conducted in different environmental contexts and with distinct organisms will be valuable to test the generality of our results.

The demonstration of large shifts in the distribution of mutational effects as a result of altered mutation bias has important implications for evolutionary dynamics. For instance, the observation that *E. coli* continues to adapt for 50,000 generations of laboratory evolution [42] begs the question of whether the distribution of beneficial mutations is finite. Our results demonstrating an immediate and substantial increase in f_b_ following a bias reversal suggest that WT *E. coli* does indeed have a finite and depleted set of beneficial mutations. Whether this is broadly true for most natural populations and species remains to be tested. Regardless, our results suggest that changes in mutation spectra across environments or populations – observed in diverse taxa [43–48] — could lead to distinct DFEs in each case, influencing evolutionary dynamics. A second important implication is regarding the evolution of mutators, which have dysfunctional DNA repair and high mutation rates that can facilitate rapid sampling of beneficial mutations [49] . Most mutators also have distinct mutation biases (e.g., except the WT, all strains used in this study are mutators), but the evolutionary implications of such bias shifts in mutators have been only rarely considered [36] . We suggest that the large DFE shifts that we observe – with concomitant changes in the beneficial mutation supply and deleterious load of mutators – can alter the rate and nature of adaptation in new environments. Accounting for these DFE shifts is thus critical to allow more accurate predictions of the fates of mutators with altered mutation biases in populations under selection. Indeed, a recent analytical and simulation study predicts that right-shifted DFEs driven by mutation bias shifts can significantly enhance the ability of mutator strains to invade non-mutator populations [12]. Thus, some mutators could gain an additional advantage – over and above the effect of high mutation rate – if their mutation bias opposes that of the ancestor. Mutators are observed frequently in natural microbial populations as well as in clinical settings, where they are often associated with drug resistance [50,51], highlighting the need to further understand the effect of mutation bias changes in mutators. More generally, our work highlights several open questions: how often is the distribution of beneficial mutations limited, how often are bias reversals observed in nature, and how often are bias shifts adaptive? Testing the impacts of mutation bias shifts on evolutionary dynamics is thus a promising direction for future research.

## Supporting information

Supplementary datasheet S1

Supplementary datasheet S2

Supplementary datasheet S3

## ACKNOWLEDGEMENTS

We thank Pratibha Sanjenbam and Lindi Wahl, and three reviewers (including Thomas Bataillon) for critical comments on the manuscript; Lindi Wahl for discussion; Kushan Lahiri for laboratory assistance; and the NGS facility for sequencing. We acknowledge funding and support from the DBT/Wellcome Trust India Alliance (Grant no. IA/S/23/2/506989 to DA), the National Centre for Biological Sciences (NCBS–TIFR) and the Department of Atomic Energy, Government of India (Project Identification No. RTI 4006), and the University Grants Commission, India (fellowship number 211610044747 to SP).

## AUTHOR CONTRIBUTIONS

MS and DA conceived of the project and designed the work; DA acquired funding; MS and SP conducted experiments and analyzed data; MS, SP and DA interpreted the results; DA wrote the manuscript with input from all authors.

