## Supplementary Materials for "Mutation bias alters the distribution of fitness effects of mutations"

### SUPPLEMENTARY FIGURES

**Figure S1. Recall rate of known background mutations.** We tested whether mutations in the background of the MG1655 clone used to construct all mutator ancestors were recovered in all evolved clones, as expected if sequencing was perfectly accurate. Violin plots show the frequency of two background mutations in our WT ancestor (compared to the NCBI reference sequence NC_000913.2) in all resequenced MA-evolved clones (A) A G→A mutation at position 2854011, and (B) An insertion of CG at position 4296830, in evolved mutator MA lines. Values under each violin are the median of the distribution.

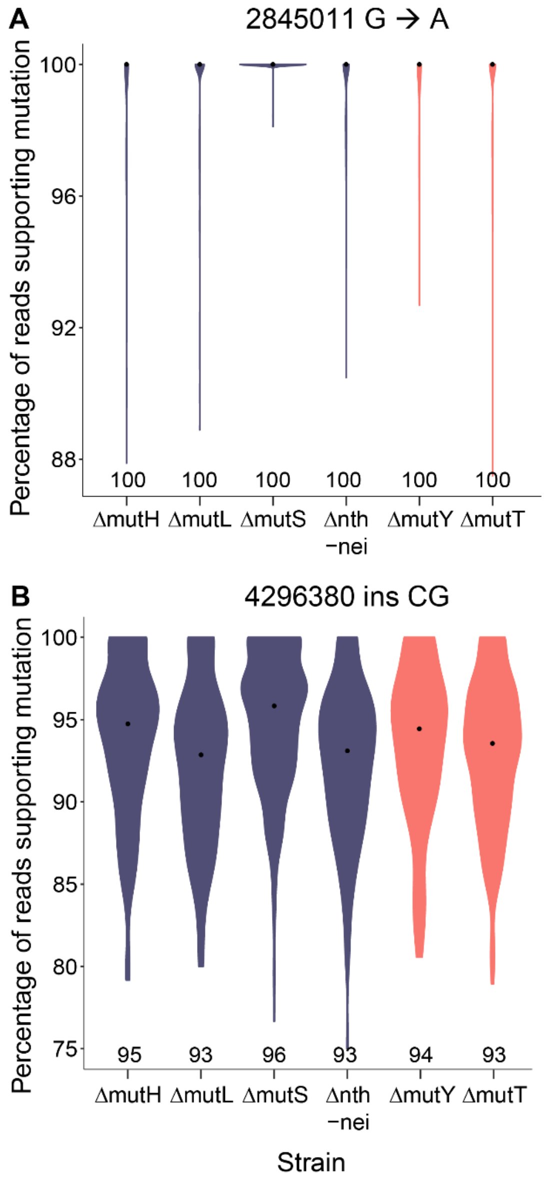

**Figure S2. The observed number of mutations per MA line is Poisson-distributed.** In each panel, open circles represent the expected number of mutations per MA line, assuming a Poisson distribution with λ=mean number of mutations observed per MA line. Filled triangles show the observed number of mutations per MA line. Results of a goodness-of-fit chi-squared test comparing observed vs. expected distributions are given in each panel. When different MA blocks differed in the number of generations evolved (in the case of WT and ΔmutY), and therefore had significantly different mean numbers of mutations per MA line across blocks, we analysed blocks separately.

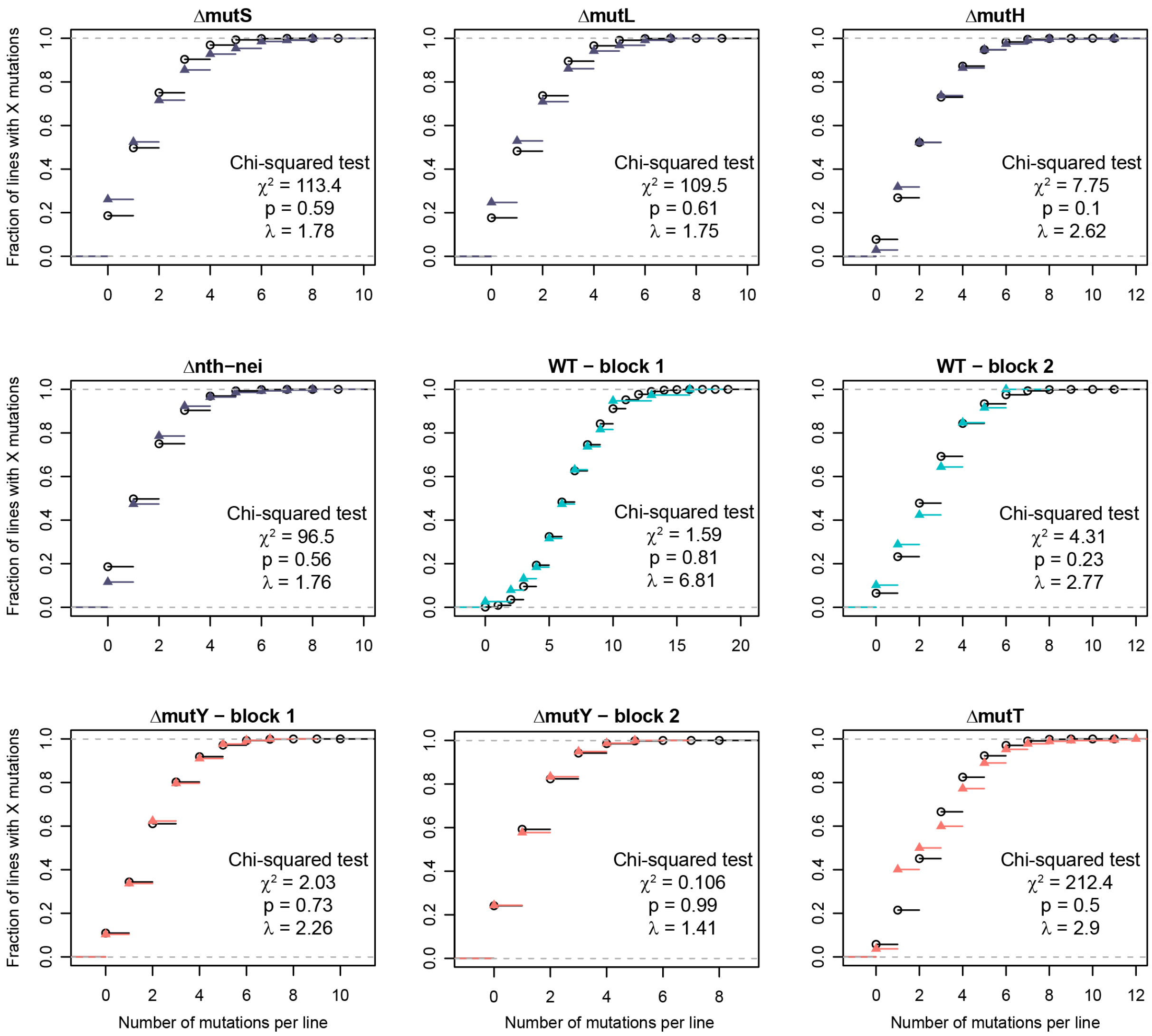

**Figure S3. Allele frequencies of mutations called in single-mutation MA lines.** Histograms show allele frequencies of mutations in MA lines included in the single mutation DFEs. In each panel, data are pooled for all MA lines included in the DFE measurements for that strain (number of lines is given in parentheses; in these MA lines, we recovered only one mutation of >80% frequency). Grey bars represent mutations segregating in MA lines at lower frequencies (<80%) and colored bars represent mutations at >80% frequency.

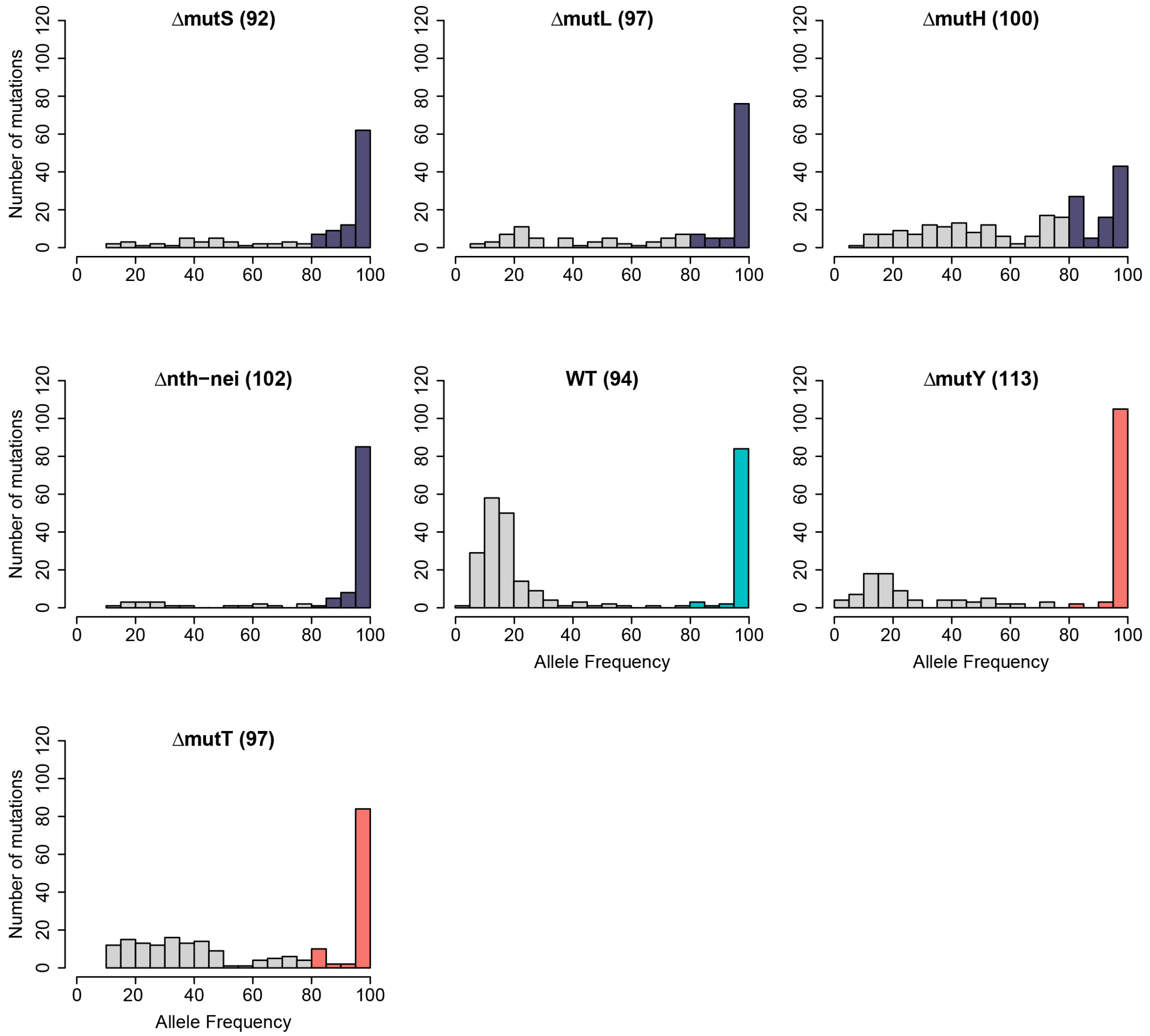

**Figure S4. Fitness measurements performed by different experimenters across years are strongly correlated.** The plot shows a fitted linear regression (dashed line) and associated statistics of the relationship between fitness measurements in Glucose conducted in 96-well plates by two different experimenters in two different years, for a set of 12 WT MA clones carrying single mutations.

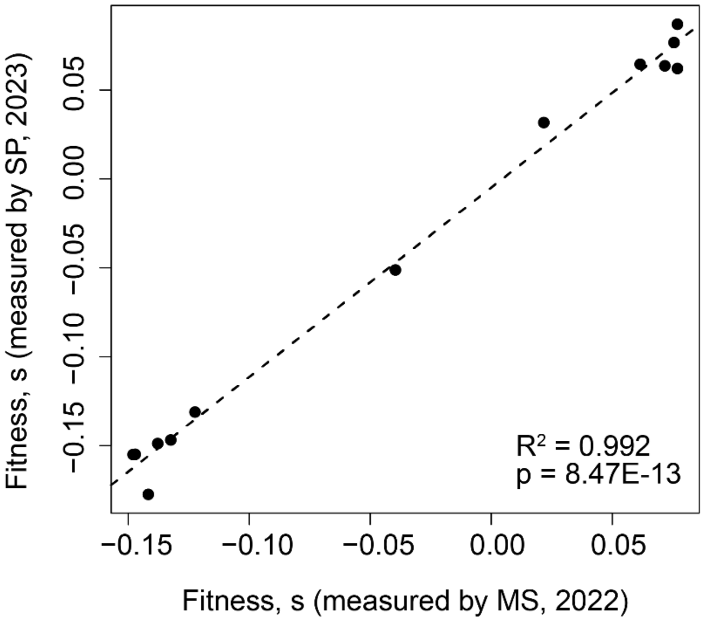


**Figure S5. Relative growth rates of single-mutation WT clones are consistent across measurements in 96-well and 48-well plates.** Each panel shows the fitted linear regression (dashed line) and associated statistics of the relationship between relative growth rates of 80 WT clones carrying single mutations obtained in 48-well plates (data from [1]) and 96-well plates (this study), in LB and Glucose.

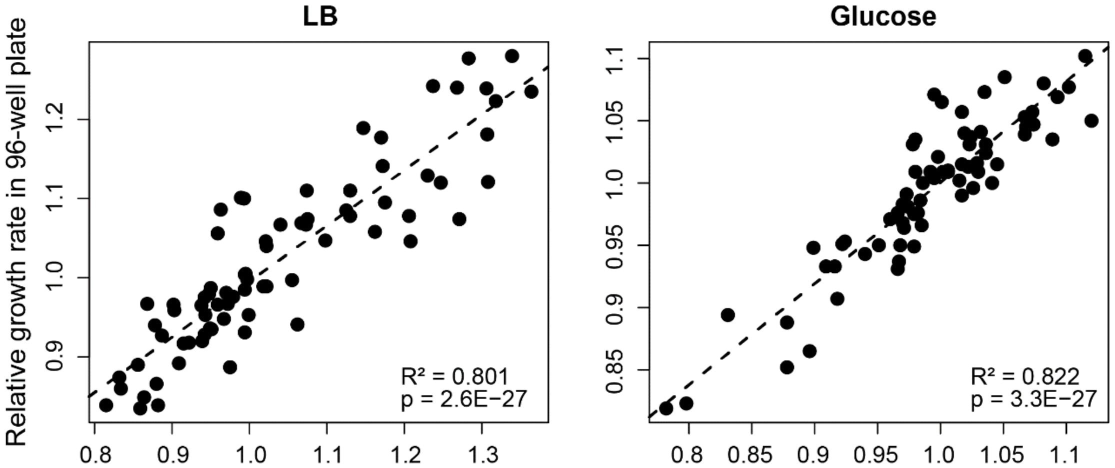

**Figure S6. Growth rates of single-mutation MA clones are strongly correlated across growth cycles.** Heritable, “real” mutations identified during resequencing should have consistent effects across successive growth cycles. The plot shows growth rates of 14 WT MA clones, each carrying a single mutation relative to the ancestor, in glucose (mean ± standard error). We show growth rates in the first 16-hour growth after reviving from frozen glycerol stocks (x-axis) vs. a second 16-hour growth cycle initiated using cultures from the first growth cycle (y-axis). The dashed line and associated statistics) represent the linear regression fit.

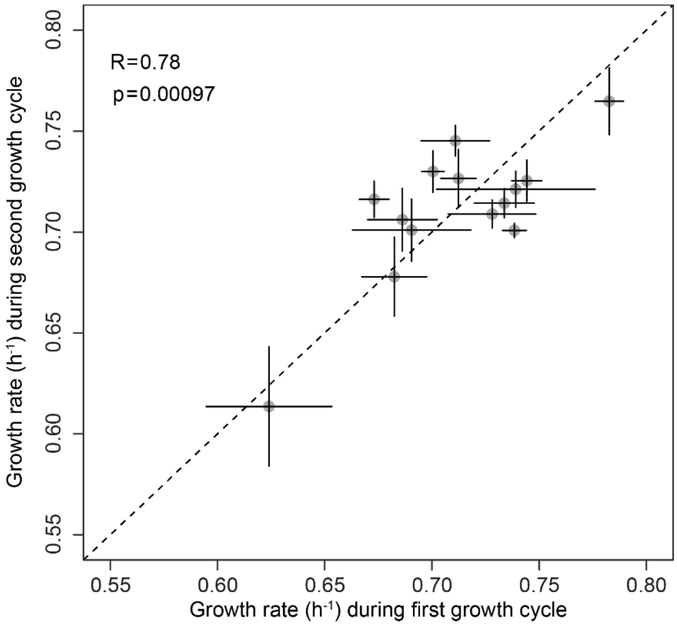

**Figure S7. Raw and selection bias-corrected DFEs of all strains in LB.** Raw (open bars) and corrected DFEs (filled bars) of single mutations in each strain's MA-accumulated mutations tested in LB. Corrected DFEs are colored as in Figure 2. Grey areas indicate neutral mutations ($s$=0±0.05 to account for experimental measurement error).

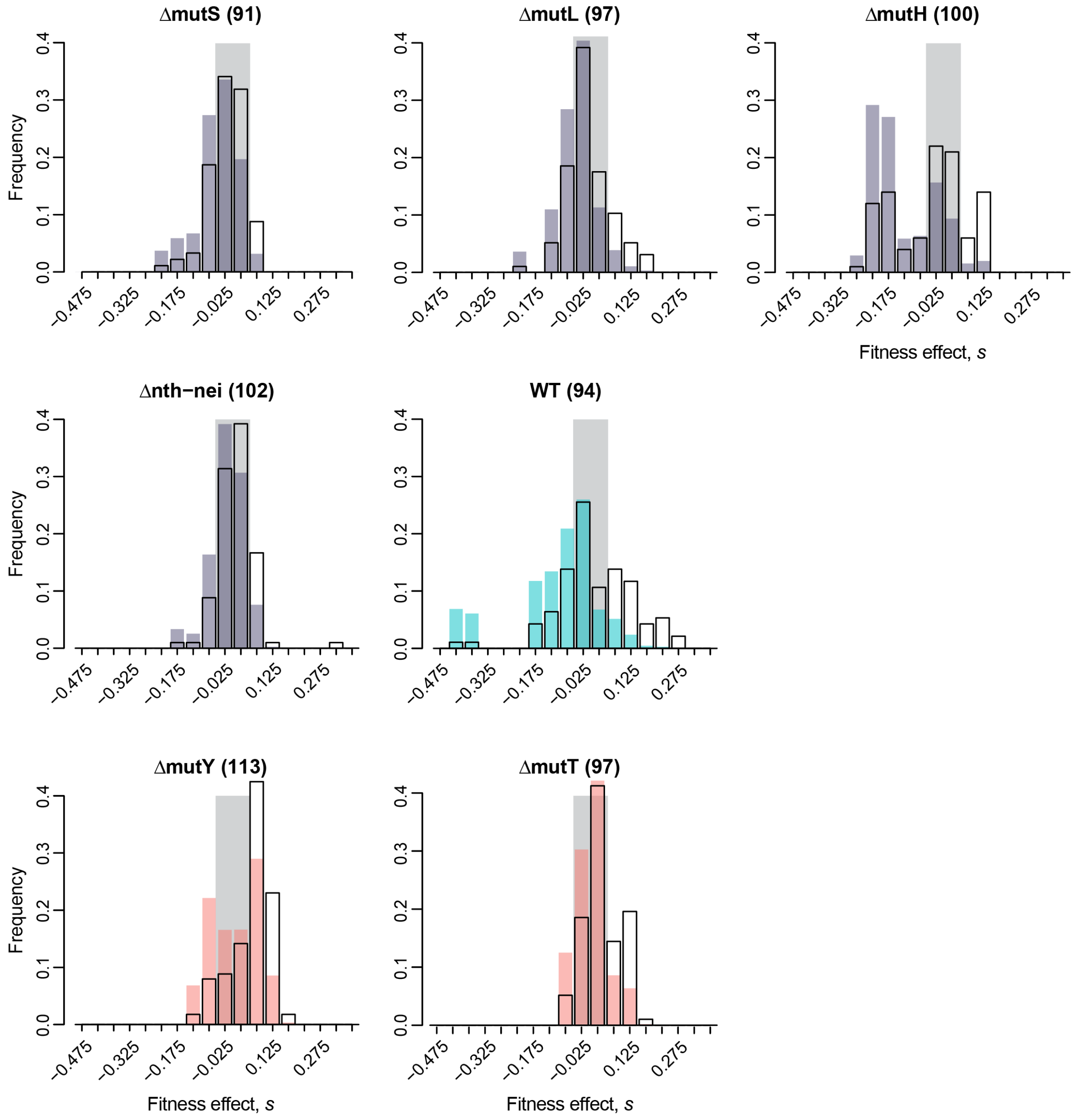

**Figure S8. Raw and selection bias-corrected DFEs of all strains in Glucose.** Raw (open bars) and corrected DFEs (filled bars) of single mutations in each strain's MA-accumulated mutations. Corrected DFEs are colored as in Figure 2. Grey areas indicate neutral mutations (*s*=0±0.025 to account for experimental measurement error).

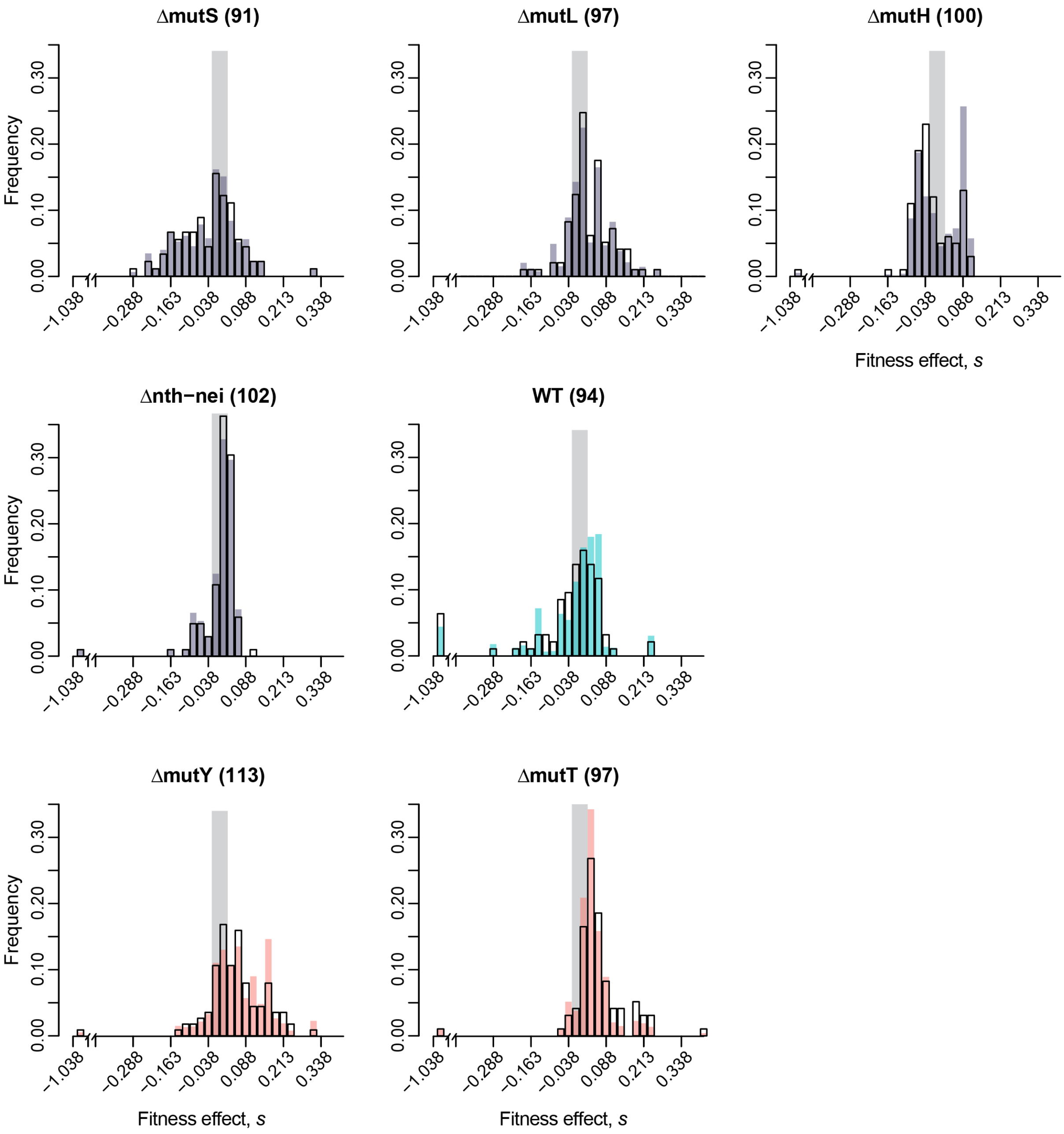

**Figure S9. Effect of stringent filtering for single mutation calling on DFEs.** The fraction of beneficial, neutral, and deleterious mutations for DFEs constructed from MA-evolved clones filtered based on the presence and frequency of secondary mutations. We applied three sets of filters to clones from each strain, comparing each DFE (after correcting for selection bias during MA) with the original ("current") DFE reported in Figure 4 (1): clones with exactly one mutation and no detectable secondary mutation, even at low frequency (2); clones with a secondary mutation at less than 20% allele frequency (3); and clones with exactly two mutations at any frequency (4). The proportion of beneficial and deleterious mutations is given in each bar. In all cases, Chi-square tests comparing each filtered set of clones with the current DFE, with Benjamini-Hochberg correction for multiple comparisons, showed a lack of significant differences in each case (p>0.05).

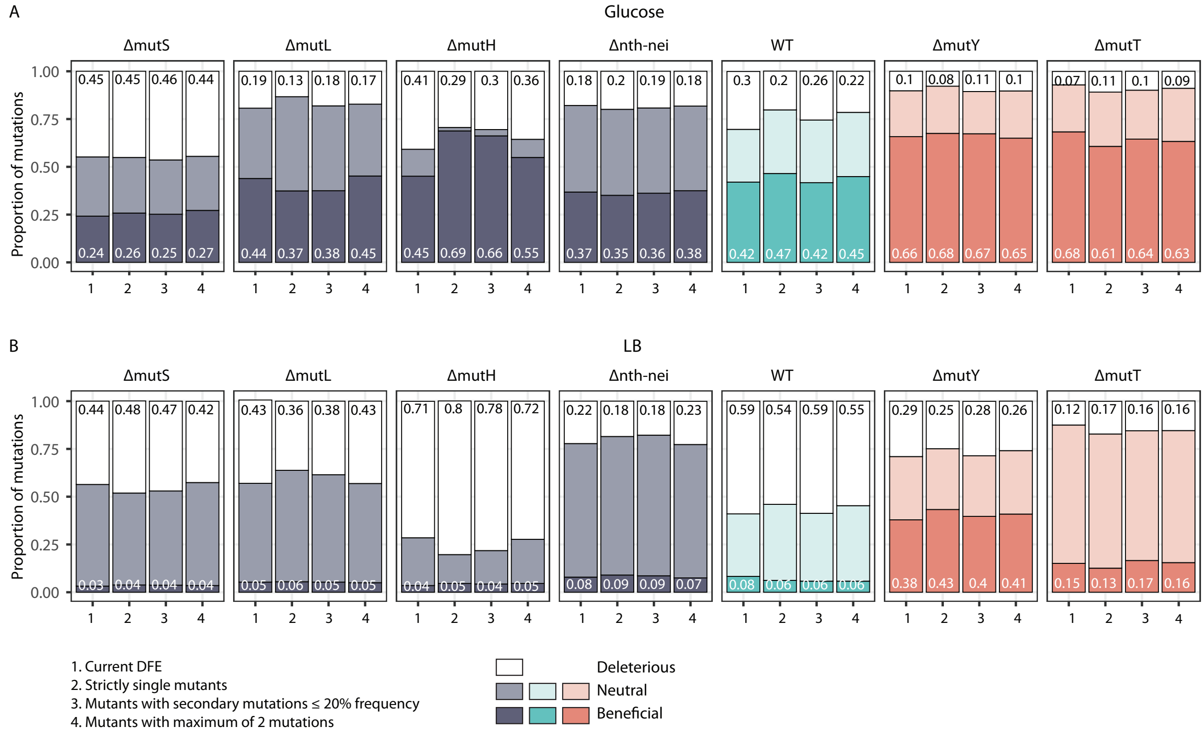

**Figure S10. Effect of reduced sample size on DFEs.** The stringent filtering described in Figure S9 reduced the number of mutants used to construct each DFE. To test the effect of reduced sample size, we subsampled the current DFE (1) reported in Figure 4 with the respective sample size of each filtered category of clones shown in Figure S9. Plots show the results from 100 iterations, with 95% confidence intervals indicated for the beneficial fraction. The sample size (number of clones) is indicated in each bar. Chi-square tests comparing each filtered set of clones with the current DFE, with Benjamini-Hochberg correction for multiple comparisons, showed a lack of significant differences (p>0.05).

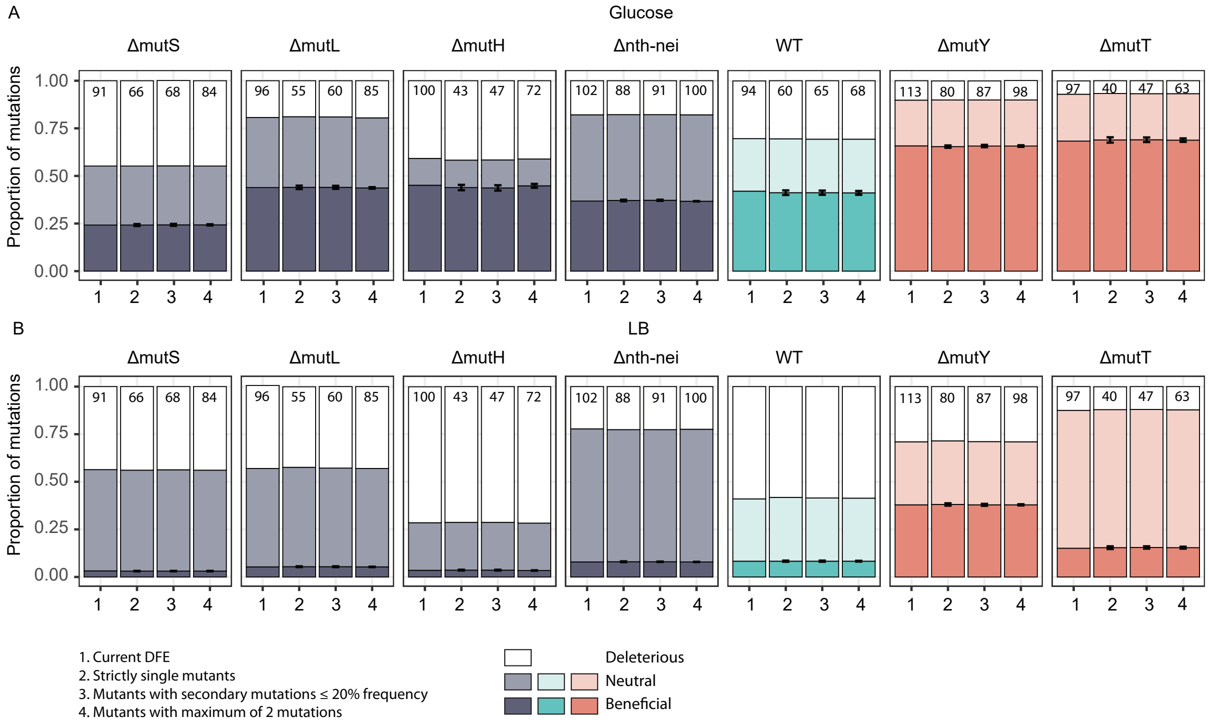

**Figure S11. The relationship between Tv bias and the fraction of beneficial ($f_b$) and deleterious mutations ($f_d$) available to strains.** Each panel shows the linear regression fit (dashed line) and associated statistics of the relationship between Tv bias and (A, B) $f_b$ or (C, D) $f_d$, in LB (left panels) and glucose (right panels). $f_b$ and $f_d$ values are given in Figure 3A and Tv bias values are given in Table 1. Points are colored as indicated in Figure 2. Error bars represent 95% confidence intervals around the mean.

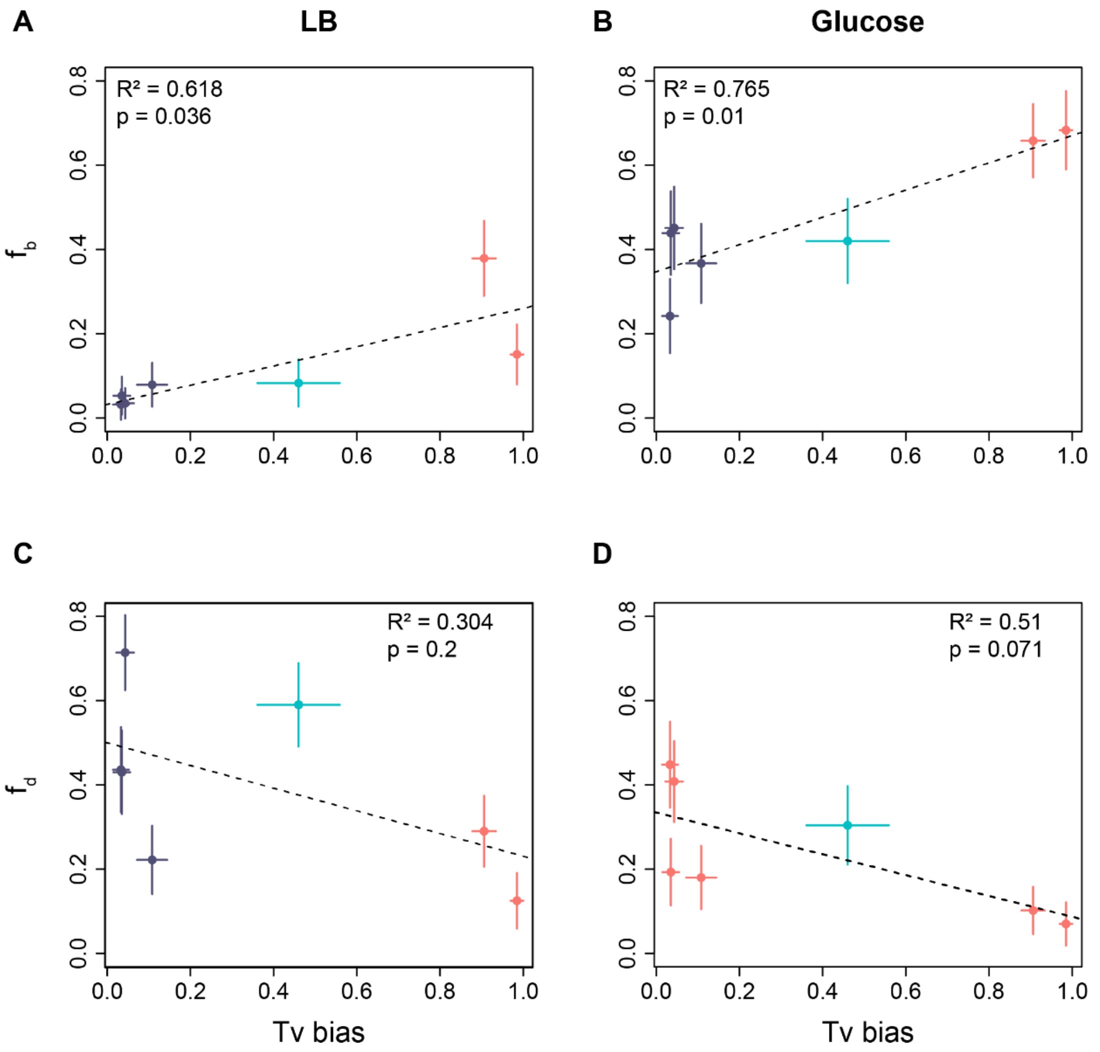

**Figure S12. The DFE alters the beneficial mutation supply and deleterious load across mutators.** Plots show the (A, B) beneficial supply and (C, D) deleterious load experienced by the different mutators as a function of mutation rate, in LB (left panels) and glucose (right panels). Strains are colored as in Figure 1 (purple: Ts-biased strains, teal: WT, pink: Tv-biased strains). Filled circles represent supply or load calculated using the $f_b$ values obtained from the observed DFE for each mutator (Figure 4A; $S_b$ and $L_d$ values shown in Tables S4 and S5 respectively); open circles represent supply or load calculated assuming $f_b$ values derived from the WT DFE (i.e., if all strains had the same DFE; $S_{b(WT\ DFE)}$ and $L_{d(WT\ DFE)}$ in Tables S4 and S5 respectively). For each strain, mutation rates used for the calculations are given in Table 1. Calculations of beneficial supply and deleterious load are shown in Tables S4 and S5. Dashed lines represent the best fit linear regression for open circles (i.e., supply or load as a function of mutation rate, assuming identical DFEs).

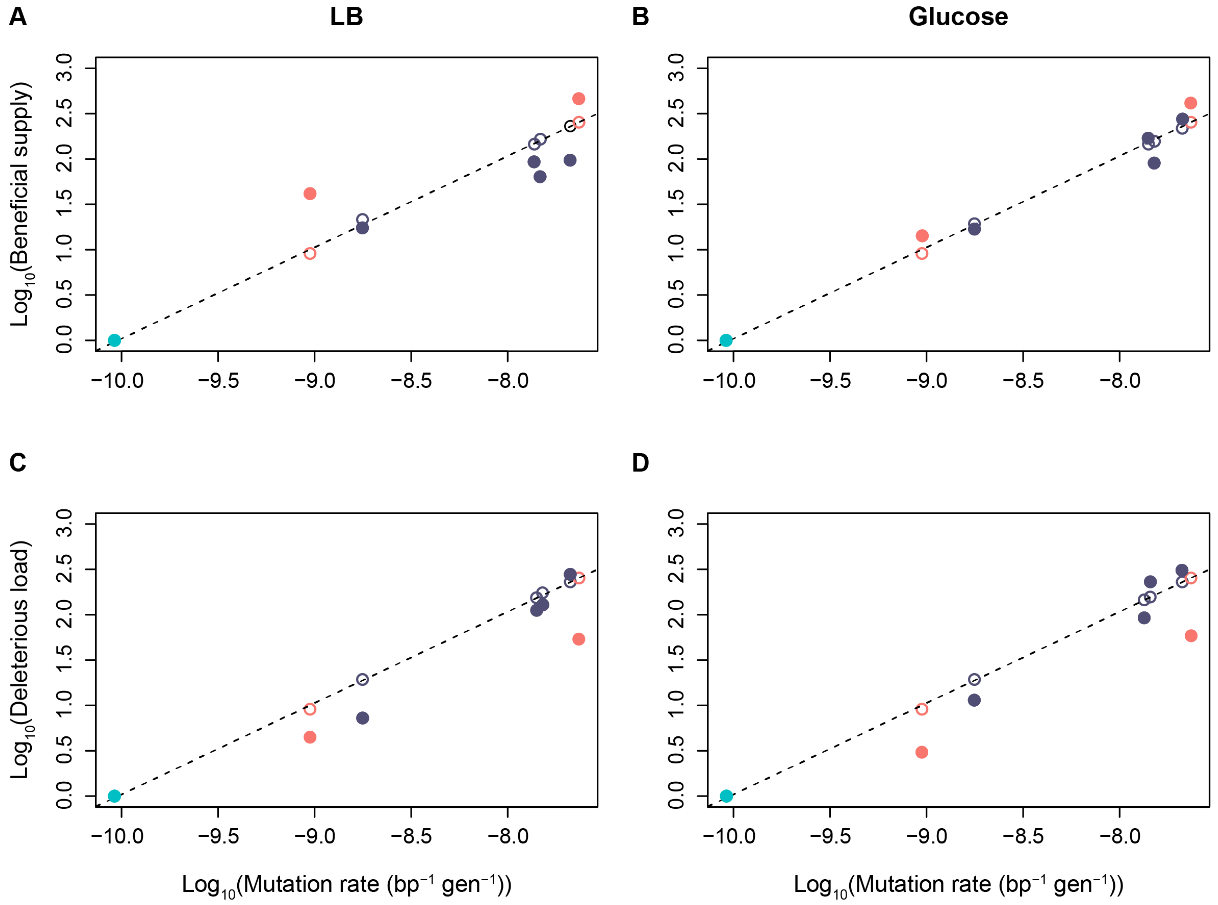

**Figure S13. Fitness effects of aspect of the mutation spectrum other than Ts/Tv.** Fitness effects of (A) BPS vs. Indel mutations, (B) Coding vs. non-coding mutations, and (C) Synonymous vs. non-synonymous mutations. In each plot, data are pooled across all strains; sample sizes (total number of single mutations tested) are shown in the LB (left) panels. When differences are significant (Wilcoxon's rank-sum tests), P values are given in the appropriate panel.

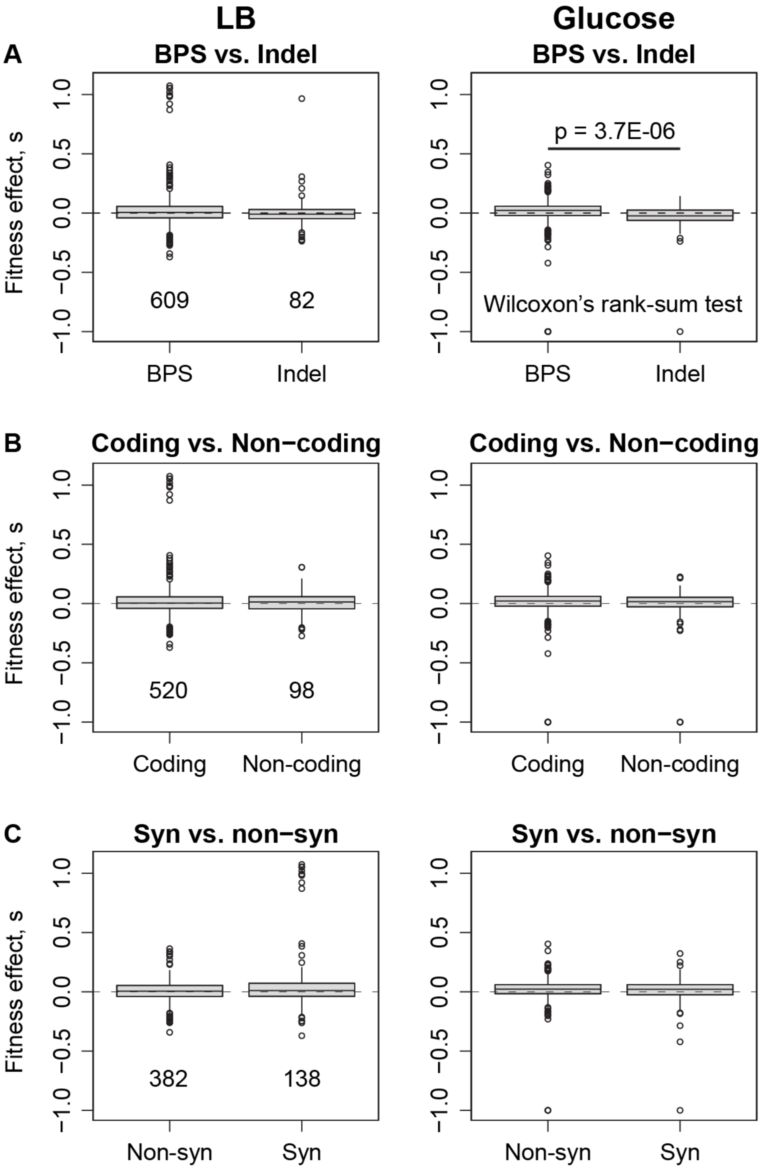

**Figure S14. The fraction of new beneficial mutations ($f_b$) available to strains does not vary with ancestral fitness.** Plots show the relationship between $f_b$ and mean ancestral growth rates in LB (left) and glucose (right). Horizontal error bars represent variation in ancestral growth rates (mean ± SE) and vertical error bars represent uncertainty in $f_b$ estimates ($f_b$ ± 95% CI). The $R^2$ and p values from a linear regression of $f_b$ ~ mean ancestral growth rate are shown in each panel.

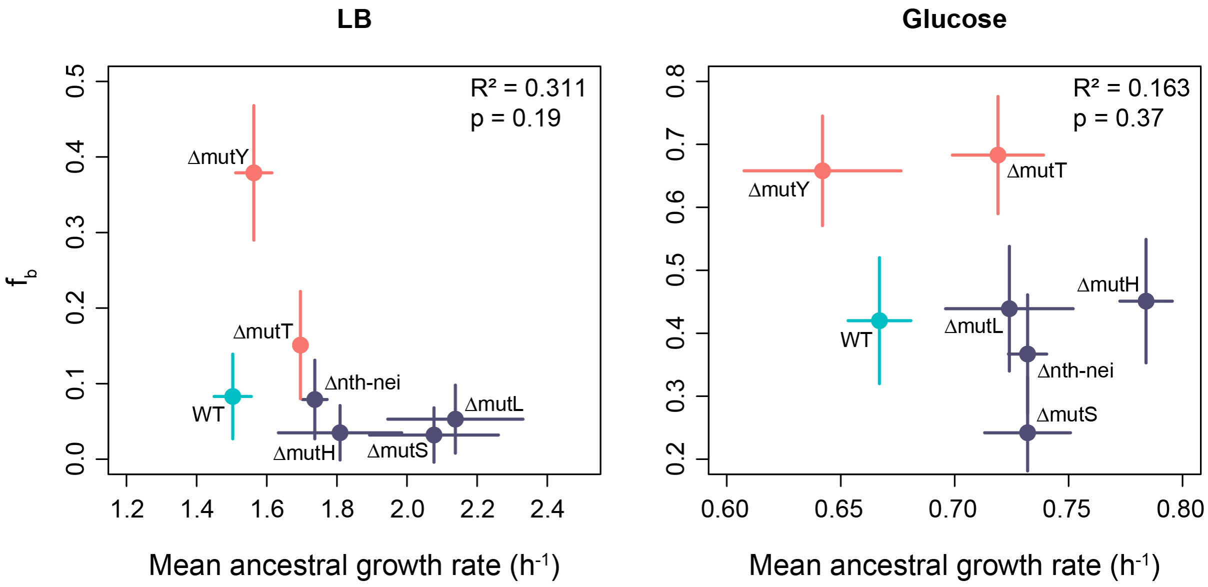

**Figure S15. Mutational effects are not associated with gene GC content.** Histograms show the distribution of gene GC content in (A) *E. coli* K-12 MG1655 genome and (B) genes with mutations in our dataset. Vertical black lines show medians. Boxplots show fitness effects of AT→GC vs. GC→AT mutations in (C, E) low GC content genes (i.e., GC content less than the genome-wide median GC) and (D, F) high GC content genes (i.e., GC content greater than the genome-wide median GC) in (C–D) LB and (E–F) Glucose. Mutational effects were not significantly different across any of the categories shown in these plots (Wilcoxon's rank-sum tests, $p > 0.05$).

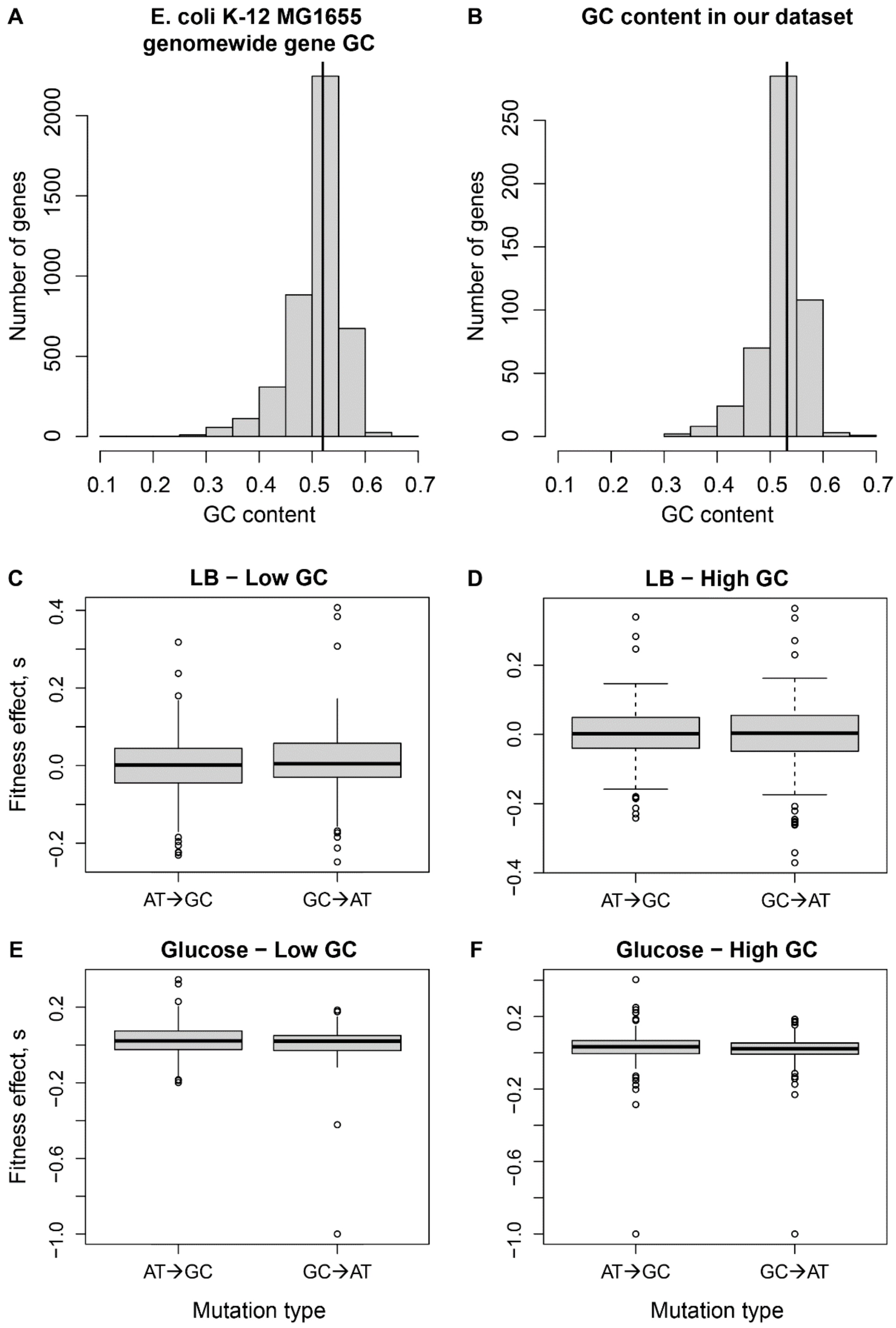

### SUPPLEMENTARY TABLES

#### Table S1: Summary of sequencing methods used in this study, and outcomes.

| Strain | Block | Type | Lines with good sequence (of X evolved)[1] | Days evolved | Lines with one mutation | Library preparation method | Sequencing platform | Mean sequencing depth (range) | Ref |
|---|---|---|---|---|---|---|---|---|---|
| ΔmutS | 1 | Anc | 1 | 0 | NA | α | A | 114x | This study |
| | | Evo | 295 (300) | 1 | 78 | | Lines 1-48: A; Lines 49-300: B | 54x (19x - 91x) | |
| | 2 | Anc | 1 | 0 | NA | | B | 43x | |
| | | Evo | 50 (50) | 1 | 14 | | B | 49x (19x - 67x) | |
| ΔmutL | 1 | Anc | 1 | 0 | NA | α | A | 88x | This study |
| | | Evo | 294 (300) | 1 | 82 | | Lines 1-48: A; Lines 49-192: B; Lines 193-300: A | 53x (18x - 126x) | |
| | 2 | Anc | 1 | 0 | NA | | A | 89x | |
| | | Evo | 50 (50) | 1 | 15 | | A | 80x (50x - 132x) | |
| ΔmutH | 1 | Anc | 1 | 0 | NA | α | A | 118x | This study |
| | | Evo | 297 (300) | 1 | 83 | | Lines 1-48: A; Lines 49-300: B | 52x (19x - 104x) | |
| | 2 | Anc | 1 | 0 | NA | | A | 66x | |
| | | Evo | 49 (50) | 1 | 17 | | A | 79x (29x - 138x) | |
| Δnth-nei | 1 | Anc | 1 | 0 | NA | α | A | 89x | This study |
| | | Evo | 80 (80) | 8 | 33 | | Lines 1-76: B; Lines 77-80: A | 61x (20x - 128x) | |
| | 2 | Anc | NA[a] | NA | NA | α | NA | NA | |
| | | Evo | 205 (220) | 8 | 69 | | A | 63x (22x - 197x) | |
| WT | 1 | Anc | 1 | 0 | 0 | β | C | 55x | [1,2] |
| | | Evo | 38 (38) | 300 | 80[b] | | A | 103x (19x – 220x) | |
| | 2 | Anc | NA[c] | NA | NA | α | NA | NA | This study |
| | | Evo | 58 (60) | 85 | 14 | | B | 64x (42x - 88x) | |
| ΔmutY | 1 | Anc | 1 | 0 | 0 | β | C | 37x | [2] |
| | | Evo | 299 (300) | 12 | 79 | | B | 82x (28x - 178x) | |
| | 2 | Anc | NA[d] | NA | NA | β | NA | NA | This study |
| | | Evo | 77 (80) | 5[e] | 26 | | B | 42x | |

| | | | | | | | | | |
|---|---|---|---|---|---|---|---|---|---|
| | | | | | | | | (21x - 61x) | |
| | 3[f] | Anc | 1 | 0 | 0 | | A | 48x | |
| | | Evo | 50 (50) | 1 | 8 | | A | 74x (47x - 109x) | |
| ΔmutT | 1 | Anc | 1 | 0 | 0 | α | A | 59x | This study |
| | | Evo | 271 (300) | 1 | 97 | | Lines 1-128: A; Lines 129-214: B; Lines 215-242: A; Lines 265-300: B; Lines 243-264: not sequenced | 55x (20x - 113x) | |

1: Lines that were not successfully sequenced (i.e., where we did not obtain sufficient good quality sequencing data or where sequencing failed entirely) are reported here, but were excluded from further analyses.
α: Illumina DNA Library preparation kit
β: Illumina Nextera XT DNA library preparation kit
A: HiSeq 2500 2x100bp PE
B: HiSeq 2500 2x125bp PE
C: MiSeq 2x250bp PE
[a, c, d] This experimental block of MA was started from the same ancestral clone as Block 1, for the respective strain.
[b] We sequenced 6 timepoints from each MA line, identifying 33 clones that had a single mutation and 29, 13 and 5 clones carrying 2, 3 or 4 mutations respectively, compared to the original ancestor. Given the low number of first-step mutants, we included the 2-, 3- and 4- step mutation clones in our dataset, but calculated their fitness relative to the immediate mutational ancestor (with the known mutation from the previously sequenced timepoint). We thus obtained a total of 80 single-mutational-step clones from 38 MA lines; see Sane et al., Evolution, 2018 for more details.
[e] This block was evolved for fewer days than Block 1, because sequencing data from Block 1 led us to revise the estimated mutation rate, allowing us to shorten the expected time to a single mutation.
[f] This block was supposed to serve as Block 2 for ΔmutT and the lines were therefore evolved for a single day. However, by mistake we used the ΔmutY ancestor to found the MA lines, so it was included in our dataset as Block 3 of ΔmutY instead of Block 2 of ΔmutT.

**Table S2: Output of chi-square tests comparing the proportion of beneficial, neutral, and deleterious mutations across strains in LB.** Values in bold highlight significant differences. Benjamini-Hochberg corrections for multiple comparisons were performed across all tests.

| Comparison | Chi-sq. statistic | P (Benjamini-Hochberg corrected) |
|---|---|---|
| ∆mutS – ∆mutL | 0.12 | 8.44E-01 |
| ∆mutS – ∆mutH | 0.00 | 1.00E+00 |
| ∆mutS – ∆nth-nei | 1.21 | 4.09E-01 |
| ∆mutS – WT | 1.38 | 4.09E-01 |
| ∆mutS – ∆mutY | 33.03 | **9.52E-08** |
| ∆mutS – ∆mutT | 6.59 | **2.75E-02** |
| ∆mutL – ∆mutH | 0.06 | 8.94E-01 |
| ∆mutL – ∆nth-nei | 0.20 | 8.05E-01 |
| ∆mutL – WT | 0.29 | 7.73E-01 |
| ∆mutL – ∆mutY | 29.74 | **3.46E-07** |
| ∆mutL – ∆mutT | 4.13 | **4.20E-02** |
| ∆mutH – ∆nth-nei | 1.03 | 4.33E-01 |
| ∆mutH – WT | 1.20 | 4.09E-01 |
| ∆mutH – ∆mutY | 34.79 | **7.73E-08** |
| ∆mutH – ∆mutT | 6.55 | **2.75E-02** |
| ∆nth-nei – WT | 0.00 | 1.00E+00 |
| ∆nth-nei – ∆mutY | 25.14 | **2.79E-06** |
| ∆nth-nei – ∆mutT | 1.92 | 3.46E-01 |
| WT – ∆mutY | 22.82 | **7.48E-06** |
| WT – ∆mutT | 1.55 | **4.06E-01** |
| ∆mutY – ∆mutT | 12.49 | **1.43E-03** |

**Table S3: Output of chi-square tests comparing the proportion of beneficial, neutral, and deleterious mutations across strains in glucose.** Values in bold highlight significant differences. Benjamini-Hochberg corrections for multiple comparisons were performed across all tests.

| Comparison | Chi-sq. statistic | P (Benjamini-Hochberg corrected) |
|---|---|---|
| ΔmutS – ΔmutL | 7.19 | **1.28E-02** |
| ΔmutS – ΔmutH | 8.22 | **7.9E-03** |
| ΔmutS – Δnth-nei | 2.96 | 1.28E-01 |
| ΔmutS – WT | 5.82 | **2.55E-02** |
| ΔmutS – ΔmutY | 33.28 | **8.35E-08** |
| ΔmutS – ΔmutT | 34.93 | **7.61E-08** |
| ΔmutL – ΔmutH | 17.48 | **3.74E-04** |
| ΔmutL – Δnth-nei | 1.55 | 4.85E-01 |
| ΔmutL – WT | 3.61 | 1.86E-01 |
| ΔmutL – ΔmutY | 10.37 | **8.41E-03** |
| ΔmutL – ΔmutT | 13.09 | **2.75E-03** |
| ΔmutH – Δnth-nei | 26.15 | **8.80E-06** |
| ΔmutH – WT | 5.85 | 6.63E-02 |
| ΔmutH – ΔmutY | 26.95 | **7.38E-06** |
| ΔmutH – ΔmutT | 30.76 | **1.47E-06** |
| Δnth-nei – WT | 7.57 | **2.98E-02** |
| Δnth-nei – ΔmutY | 18.15 | **3.00E-04** |
| Δnth-nei – ΔmutT | 20.29 | **1.24E-04** |
| WT – ΔmutY | 16.31 | **6.02E-04** |
| WT – ΔmutT | 20.18 | **1.24E-04** |
| ΔmutY – ΔmutT | 0.66 | 7.19E-01 |

**Table S4. Beneficial supply calculations for all strains.** Supply of beneficial mutations ($S_b$) is calculated for each strain in both environments (LB and Glucose) using the empirically estimated $f_b$ values (Figure 4A), whole genome mutation rates (μ, Table 1), and genome size (4641652 bp) as: $S_b = f_b \times \mu \times$ genome size. Beneficial supply assuming a WT DFE, $S_{b(WT\ DFE)}$, is calculated as $f_{b(WT)} \times \mu \times$ genome size. $S_b$ and $S_{b(WT\ DFE)}$ relative to WT are reported in the two rightmost columns. Confidence intervals were calculated as 1.96 x (standard deviation of $S_b$).

| Strain | Env | $f_b$ | μ | $S_b$ | $S_{b(WT\ DFE)}$ | $S_b/S_{b(WT)}$ | $S_{b(WT\ DFE)}/S_{b(WT)}$ |
|---|---|---|---|---|---|---|---|
| ΔmutS | LB | 0.032 ±0.04 | 1.44E-08 | 2.14E-03 ±2.27E-03 | 5.54E-03 | 60.35 | 156.53 |
| ΔmutL | LB | 0.053 ±0.04 | 1.41E-08 | 3.47E-03 ±2.62E-03 | 5.44E-03 | 98.12 | 153.67 |
| ΔmutH | LB | 0.035 ±0.04 | 2.11E-08 | 3.43E-03 ±2.23E-03 | 8.14E-03 | 97.01 | 230.06 |
| Δnth_nei | LB | 0.079 ±0.05 | 1.77E-09 | 6.50E-04 ±3.06E-04 | 6.83E-04 | 18.37 | 19.30 |
| WT | LB | 0.083 ±0.06 | 9.18E-11 | 3.54E-05 ±3.70E-05 | 3.54E-05 | 1.00 | 1.00 |
| ΔmutY | LB | 0.379 ±0.09 | 8.34E-10 | 1.47E-03 ±7.06E-04 | 3.21E-04 | 41.48 | 9.08 |
| ΔmutT | LB | 0.151 ±0.07 | 2.34E-08 | 1.64E-02 ±5.47E-03 | 9.00E-03 | 462.63 | 254.29 |
| ΔmutS | Glu | 0.242 ±0.09 | 1.44E-08 | 1.61E-02 ±5.52E-03 | 2.80E-02 | 90.19 | 156.53 |
| ΔmutL | Glu | 0.393 ±0.10 | 1.41E-08 | 2.57E-02 ±5.71E-03 | 2.75E-02 | 143.79 | 153.67 |
| ΔmutH | Glu | 0.451 ±0.10 | 2.11E-08 | 4.42E-02 ±6.04E-03 | 4.12E-02 | 247.04 | 230.06 |
| Δnth_nei | Glu | 0.367 ±0.09 | 1.77E-09 | 3.02E-03 ±5.46E-04 | 3.46E-03 | 16.87 | 19.30 |
| WT | Glu | 0.42 ±0.10 | 9.18E-11 | 1.79E-04 ±6.62E-05 | 1.79E-04 | 1.00 | 1.00 |
| ΔmutY | Glu | 0.658 ±0.09 | 8.34E-10 | 2.55E-03 ±6.91E-04 | 1.63E-03 | 14.23 | 9.08 |
| ΔmutT | Glu | 0.683 ±0.09 | 2.34E-08 | 7.40E-02 ±7.10E-03 | 4.55E-02 | 413.53 | 254.29 |

**Table S5. Deleterious load calculations for all strains.** Deleterious load ($L_d$) is calculated for each strain in both environments (LB and Glucose) using the empirically estimated $f_d$ values (Figure 4A), whole genome mutation rates (μ) (Table 1), and genome size (4641652 bp) as: $L_d = f_d \times \mu \times$ genome size. Deleterious load assuming a WT DFE, $L_{d(WT\ DFE)}$, is calculated as $f_{d(WT)} \times \mu \times$ genome size. $L_d$ and $L_{d(WT\ DFE)}$ relative to WT are reported in the two rightmost columns. Confidence intervals were calculated as 1.96 x (standard deviation of $L_d$).

| Strain | Env | $f_d$ | μ | $L_d$ | $L_{d(WT\ DFE)}$ | $L_d$ /$L_{d(WT)}$ | $L_{d(WT\ DFE)}$/ $L_{d(WT)}$ |
|---|---|---|---|---|---|---|---|
| ΔmutS | LB | 0.44 ±0.10 | 1.44E-08 | 2.91E-02 ±6.39E-03 | 3.94E-02 | 115.67 | 156.53 |
| ΔmutL | LB | 0.43 ±0.10 | 1.41E-08 | 2.82E-02 ±5.79E-03 | 3.87E-02 | 111.99 | 153.67 |
| ΔmutH | LB | 0.71 ±0.09 | 2.11E-08 | 7.00E-02 ±5.48E-03 | 5.79E-02 | 278.41 | 230.06 |
| Δnth-nei | LB | 0.22 ±0.08 | 1.77E-09 | 1.83E-03 ±4.71E-04 | 4.85E-03 | 7.26 | 19.30 |
| WT | LB | 0.59 ±0.10 | 9.18E-11 | 2.52E-04 ±6.60E-05 | 2.52E-04 | 1.00 | 1.00 |
| ΔmutY | LB | 0.29 ±0.08 | 8.34E-10 | 1.12E-03 ±6.61E-04 | 2.28E-03 | 4.46 | 9.08 |
| ΔmutT | LB | 0.13 ±0.07 | 2.34E-08 | 1.36E-02 ±5.05E-03 | 6.40E-02 | 53.88 | 254.29 |
| ΔmutS | Glu | 0.45 ±0.10 | 1.44E-08 | 2.99E-02 ±6.41E-03 | 2.03E-02 | 230.67 | 156.53 |
| ΔmutL | Glu | 0.23 ±0.08 | 1.41E-08 | 1.51E-02 ±4.93E-03 | 1.99E-02 | 116.77 | 153.67 |
| ΔmutH | Glu | 0.41 ±0.10 | 2.11E-08 | 4.00E-02 ±5.96E-03 | 2.98E-02 | 308.76 | 230.06 |
| Δnth-nei | Glu | 0.18 ±0.07 | 1.77E-09 | 1.48E-03 ±4.35E-04 | 2.50E-03 | 11.43 | 19.30 |
| WT | Glu | 0.30 ±0.09 | 9.18E-11 | 1.30E-04 ±6.17E-05 | 1.30E-04 | 1.00 | 1.00 |
| ΔmutY | Glu | 0.10 ±0.06 | 8.34E-10 | 3.95E-04 ±4.41E-04 | 1.18E-03 | 3.05 | 9.08 |
| ΔmutT | Glu | 0.07 ±0.05 | 2.34E-08 | 7.59E-03 ±3.90E-03 | 3.30E-02 | 58.55 | 254.29 |

**Table S6. Fitness effects of mutations associated with other axes of mutation bias.** The table shows differences between fitness effects of different types of mutations between pairs of strains. Values in bold highlight significant differences.

| Mutation type | Comparison | P (Benjamini-Hochberg corrected) | |
|---|---|---|---|
| | | LB | Glucose |
| coding | ΔmutL – ΔmutH | 0.538 | **0.002** |
| coding | ΔmutS – ΔmutH | 0.297 | 0.134 |
| coding | ΔmutS – ΔmutL | 0.539 | **0.001** |
| coding | Δnth-nei – ΔmutH | 0.063 | 0.103 |
| coding | Δnth-nei – ΔmutL | 0.063 | 0.050 |
| coding | Δnth-nei – ΔmutS | 0.209 | **0.014** |
| coding | WT – ΔmutH | **0.009** | 0.346 |
| coding | WT – ΔmutL | 0.063 | **0.014** |
| coding | WT – ΔmutS | 0.098 | 0.106 |
| coding | WT – Δnth-nei | 0.297 | 0.346 |
| noncoding | ΔmutL – ΔmutH | 0.989 | **0.018** |
| noncoding | ΔmutS – ΔmutH | 0.795 | 0.572 |
| noncoding | ΔmutS – ΔmutL | 0.613 | 0.109 |
| noncoding | Δnth-nei – ΔmutH | 0.795 | 0.064 |
| noncoding | Δnth-nei – ΔmutL | 0.334 | 0.352 |
| noncoding | Δnth-nei – ΔmutS | **0.018** | 0.572 |
| noncoding | WT – ΔmutH | 0.519 | 0.791 |
| noncoding | WT – ΔmutL | 0.519 | **0.045** |
| noncoding | WT – ΔmutS | 0.519 | 0.747 |
| noncoding | WT – Δnth-nei | 0.989 | 0.109 |
| syn | ΔmutL – ΔmutH | 0.897 | 0.415 |
| syn | ΔmutS – ΔmutH | 0.897 | 0.534 |
| syn | ΔmutS – ΔmutL | 0.897 | 0.415 |
| syn | Δnth-nei – ΔmutH | 0.135 | 0.671 |
| syn | Δnth-nei – ΔmutL | 0.135 | 0.534 |
| syn | Δnth-nei – ΔmutS | 0.266 | 0.415 |
| syn | WT – ΔmutH | 0.093 | 0.691 |
| syn | WT – ΔmutL | 0.104 | 0.415 |
| syn | WT – ΔmutS | 0.104 | 0.453 |
| syn | WT – Δnth-nei | 0.583 | 0.560 |
| nonsyn | ΔmutL – ΔmutH | 0.552 | **0.006** |
| nonsyn | ΔmutS – ΔmutH | 0.552 | 0.256 |
| nonsyn | ΔmutS – ΔmutL | 0.552 | **0.006** |
| nonsyn | Δnth-nei – ΔmutH | 0.199 | 0.118 |
| nonsyn | Δnth-nei – ΔmutL | 0.199 | 0.118 |
| nonsyn | Δnth-nei – ΔmutS | 0.552 | 0.076 |
| nonsyn | WT – ΔmutH | 0.199 | 0.181 |
| nonsyn | WT – ΔmutL | 0.552 | 0.147 |
| nonsyn | WT – ΔmutS | 0.653 | 0.118 |
| nonsyn | WT – Δnth-nei | 0.997 | 0.838 |
| AT→GC | ΔmutL – ΔmutH | 0.973 | **0.000** |
| AT→GC | ΔmutS – ΔmutH | 0.973 | 0.695 |

| | | | |
|---|---|---|---|
| AT→GC | ΔmutS – ΔmutL | 0.973 | **0.002** |
| AT→GC | Δnth-nei – ΔmutH | 0.973 | 0.695 |
| AT→GC | Δnth-nei – ΔmutL | 0.973 | 0.292 |
| AT→GC | Δnth-nei – ΔmutS | 0.973 | 0.695 |
| AT→GC | WT – ΔmutH | 0.149 | 0.342 |
| AT→GC | WT – ΔmutL | 0.167 | 0.181 |
| AT→GC | WT – ΔmutS | 0.149 | 0.342 |
| AT→GC | WT – Δnth-nei | 0.903 | 0.712 |
| GC→AT | ΔmutL – ΔmutH | 0.667 | 0.256 |
| GC→AT | ΔmutS – ΔmutH | 0.317 | 0.256 |
| GC→AT | ΔmutS – ΔmutL | 0.685 | **0.029** |
| GC→AT | Δnth-nei – ΔmutH | 0.317 | 0.577 |
| GC→AT | Δnth-nei – ΔmutL | 0.667 | 0.512 |
| GC→AT | Δnth-nei – ΔmutS | 0.880 | **0.029** |
| GC→AT | WT – ΔmutH | 0.317 | 0.828 |
| GC→AT | WT – ΔmutL | 0.812 | 0.256 |
| GC→AT | WT – ΔmutS | 0.880 | 0.202 |
| GC→AT | WT – Δnth-nei | 0.880 | 0.256 |
